# Isolation of adipose tissue derived regenerative cells from human subcutaneous tissue with or without the use of an enzymatic reagent

**DOI:** 10.1101/485318

**Authors:** Glenn E. Winnier, Nick Valenzuela, Christopher Alt, Jennifer Peters-Hall, Joshua Kellner, Eckhard U. Alt

**Affiliations:** InGeneron, Inc., Houston, TX, USA; Heart and Vascular Institute, Department of Medicine, Tulane University Health Science Center, New Orleans, LA, USA; Sanford Health, University of South Dakota, Sioux Falls, SD, USA; Isar Klinikum Munich, Munich, Germany

## Abstract

**Background:** Freshly isolated, uncultured autologous adipose derived regenerative cells (ADRCs) have emerged as a promising tool for regenerative cell therapy. The Transpose RT system (InGeneron, Inc., Houston, TX, USA) is a system for isolating ADRCs from adipose tissue, commercially available in Europe as a CE-marked medical device and under clinical evaluation in the United States. This system makes use of the proprietary, enzymatic Matrase Reagent for isolating cells. The present study addressed the question whether the use of Matrase Reagent influences cell yield, cell viability, live cell yield, biological characteristics, physiological functions or structural properties of the ADRCs in final cell suspension.

**Methods:** Identical samples of subcutaneous adipose tissue from 12 subjects undergoing elective lipoplasty were processed either with or without the use of Matrase Reagent. Then, characteristics of the ADRCs in the respective final cell suspensions were evaluated.

**Results:** Compared to non-enzymatic isolation, enzymatic isolation resulted in approximately twelve times higher mean cell yield (i.e., numbers of viable cells/ml lipoaspirate) and approximately 16 times more colony forming units. Despite these differences, cells isolated from lipoaspirate both with and without the use of Matrase Reagent were independently able to differentiate into cells of all three germ layers.

**Discussion:** The data of this study indicate that biological characteristics, physiological functions or structural properties relevant for the intended use were not altered or induced using Matrase Reagent. A comprehensive literature review demonstrated that isolation of ADRCs from lipoaspirate using the Transpose RT system and the Matrase Reagent results in the highest viable cell yield among all published data regarding isolation of ADRCs from lipoaspirate.

## INTRODUCTION

Regenerative cell therapy, which refers to the therapeutic application of stem cells to repair diseased or injured tissue, has received increasing attention from basic scientists, clinicians and the public (e.g., [1–5]). Stem cells hold significant promise for tissue regeneration due to their innate ability to provide a renewable supply of progenitor cells that can form multiple cell types, whole tissue structures, and even organs (e.g., [1–5]).

Over the last years, fresh, uncultured, autologous adipose-derived regenerative cells (UA-ADRCs) have become highly attractive for the practice of regenerative cell therapy (e.g., [6–10]) (note that fresh, uncultured, adipose-derived regenerative cells were also named “stromal vascular fraction” (SVF) in many publications; e.g., [11–13]). This is due to the fact that UA-ADRCs have several advantages over other types of cells used in and/or under investigation for regenerative cell therapy. First of all, UA-ADRCs do not share ethical concerns nor the risk of teratoma formation reported for embryonic stem cells [14–16]. Neither do UA-ADRCs share the risk of tumorigenesis that severely limits the clinical translation of induced pluripotent stem (iPS) cells [17–19]. Second, because UA-ADRCs are autologous cells, their application does not bear the risk of HLA mismatch associated with allogeneic cells (compromised clinical outcome after application of allogeneic cells was reported in [20–22]). Third, adipose tissue typically has a significantly higher stem cell density than bone marrow (5 to 10% vs. 0.1%), and harvesting adipose tissue can be less invasive than harvesting bone marrow [23, 24]. Fourth, unlike the use of adipose derived stem cells (ASCs) which are culture-expanded from the SVF [25–27], the use of UA-ADRCs allows immediate usage at point of care [8–10]. This is combined with low safety concerns as no culturing or modification are applied. Of note, when using UA-ADRCs during a surgical procedure in an autologous and homologous way, they are not considered an advanced therapy medicinal product (ATMP) by the European Medicines Agency [28]. In contrast, expansion of ASCs in vitro may be associated with risks such as possible loss of stemness or cell transformation [29–31]. On the other hand, recent studies on culture systems and animal models indicated non-inferiority or even superiority of UA-ADRCs over ASCs in, for example, rescuing heart function after acute myocardial infarction [32], bone regeneration [33] and erectile function recovery after cavernous nerve injury [34] (see also [13]).

For obvious reasons, an optimal system for providing UA-ADRCs at point of care should be capable of isolating the highest possible number of living ADRCs from the lowest possible amount of adipose tissue in the shortest possible time, and providing the cells at the highest possible concentration in a final cell suspension.

A number of enzymatic and non-enzymatic methods were reported for isolating human ADRCs (Tables 1 and 2; c.f. also Fig 1 and [35–38]). For enzymatic methods, cell yield between 0.0 [39] and 387×10^5^ cells per ml lipoaspirate [63] were reported (Table 1) (mean, 14.6×10^5^; standard deviation, 61.5×10^5^; median, 3.1×10^5^; for 90% of the methods listed in Table 1 cell yield between 0.0 and 12.2×10^5^ cells per ml lipoaspirate was reported; Fig 1). Related cell viability data (i.e., relative number of living cells) varied between 50% [40] and 94% [23] (note that for 14 out of the 39 enzymatic methods no cell viability data were reported; Table 1).

**Table 1.** Cell yield, cell viability, number of living cells per gram lipoaspirate and other details reported in studies describing enzymatic methods for isolating ADRCs.

| R | Y | S | O/C | M/S/A | CY | V | LCY | V<sub>L</sub> | PT | FV |
| --- | --- | --- | --- | --- | --- | --- | --- | --- | --- | --- |
| [39] | 2015 | (a) | O | M | **0.0** | -- | -- | 80 | -- | -- |
| [40] | 2013 | (b) | C | S | **0.07** | 87 | 0.06 | 130 | 88 | -- |
| [40] | 2013 | (c) | C | M | **0.35** | 72 | 0.25 | 80 | 111 | -- |
| [41] | 2014 | (d) | C | A | **0.72** | 91 | 0.66 | 250 | 98 | -- |
| [42] | 2015 |  | C | A | **1.00** | >90 | 0.90 | 500 | 133 | 11 |
| [43] | 2016 | (e) | C | A | **1.01** | 84 | 0.85 | 162 | 89 | 5 |
| [42] | 2015 |  | O | M | **1.03** | >90 | 0.92 | -- | -- | -- |
| [40] | 2013 | (f) | O | M | **1.07** | 57 | 0.61 | 125 | 115 | -- |
| [42] | 2015 |  | O | M | **1.47** | -- | -- | 212 | -- | -- |
| [44] | 2012 |  | O | M | **1.60** | >90 | 1.44 | 220 | -- | -- |
| [45] | 2014 |  | O | M | **2.30** | -- | -- | -- | -- | -- |
| [40] | 2013 | (g) | C | A | **2.41** | 93 | 2.24 | 140 | 90 | -- |
| [39] | 2015 | (h) | C | A | **2.55** | -- | -- | 80 | -- | -- |
| [44] | 2012 |  | C | S | **2.60** | >90 | 2.34 | 220 | -- | 10 |
| [43] | 2016 | (i) | C | M | **2.85** | 69 | 1.96 | 102 | 71 | 7 |
| [46] | 2019 |  | O | M | **2.93** | 76 | 2.23 | 100 | 82 | -- |
| [47] | 2015 | (j) | C | A | **2.79** | 88 | 2.46 | 181 | 120 | 8 |
| [48] | 2008 | (k) | C | A | **2.95** | 87 | 2.57 | 86 | -- | -- |
| [49] | 2005 | (l) | C | A | **3.00** | 93 | 2.79 | -- | -- | -- |
| [50] | 2006 |  | O | M | **3.08** | -- | -- | -- | -- | -- |
| [51] | 2014 | (m) | C | A | **3.60** | 85 | 3.06 | -- | -- | 5 |
| [52] | 2014 | (n) | C | M | **3.60** | 83 | 2.98 | 400 | -- | -- |
| [53] | 2014 |  | O | M | **3.60** | 67 | 2.41 | -- | -- | -- |
| [54] | 2014 |  | O | M | **3.68** | 75 | 2.76 | -- | -- | -- |
| [23] | 2004 |  | O | M | **4.04** | 94 | 3.80 | -- | -- | -- |
| [55] | 2013 |  | O | M | **4.07** | -- | -- | 24 | -- | -- |
| [56] | 2013 |  | O | M | **4.80** | -- | -- | 55 | -- | -- |
| [57] | 2015 | (o) | C | M | **5.34** | -- | -- | -- | -- | -- |
| [46] | 2019 |  | O | M | **5.35** | 86 | 4.60 | 100 | 72 | -- |
| [43] | 2016 | (p) | O | M | **5.36** | 82 | 4.39 | 156 | 65 | 12 |
| [43] | 2016 | (q) | C | M | **6.25** | 50 | 3.12 | 140 | 121 | 20 |
| [58] | 2013 |  | O | M | **7.01** | 82 | 5.75 | 50 | -- | -- |
| [58] | 2013 |  | C | A | **7.02** | 81 | 5.69 | 50 | -- | -- |
| [59] | 2017 | (r) | C | M | **9.06** | -- | -- | 50 | 120 | -- |
| [57] | 2015 | (s) | C | A | **12.1** | -- | -- | -- | -- | -- |
| [60] | 2006 |  | O | M | **13.1** | -- | -- | -- | -- | -- |
| [61] | 2001 |  | O | M | **13.3** | -- | -- | 300 | -- | -- |
| [62] | 2016 | (t) | C | M | **35.0** | -- | -- | -- | 80 | 5 |
| [63] | 2016 |  | O | M | **387** | -- | -- | 80 | -- | -- |

**Table 2.** Cell yield, cell viability, number of living cells per gram lipoaspirate and other details reported in studies describing non-enzymatic methods for isolating ADRCs.

| R | Y | S | O/C | M/S/A | CY | CV | LCY | V<sub>L</sub> | PT | FV |
| --- | --- | --- | --- | --- | --- | --- | --- | --- | --- | --- |
| [39] | 2015 | (u) | O | S | **0.07** | -- | -- | 80 | -- | -- |
| [53] | 2014 |  | O | M | **0.11** | 69 | 0.08 | -- | -- | -- |
| [45] | 2014 |  | O | M | **0.12** | -- | -- | -- | -- | -- |
| [45] | 2014 |  | O | M | **0.23** | -- | -- | -- | -- | -- |
| [56] | 2013 |  | O | M | **0.25** | -- | -- | 180 | -- | -- |
| [39] | 2015 | (v) | O | M | **0.30** | -- | -- | 80 | -- | -- |
| [64] | 2014 |  | O | M | **1.25** | -- | -- | 80 | 15 | -- |
| [54] | 2014 | (i) | C | S | **1.39** | -- | -- | 200 | -- | -- |
| [65] | 2009 |  | O | M | **2.40** | -- | -- | -- | -- | -- |
| [57] | 2015 | (j) |  |  | **4.44** | -- | -- | 80 | -- | -- |

Abbreviations: R, reference; Y, year of publication; S, name of commercial system (if applicable; information is given here as provided in the corresponding references); O, open isolation system; C, closed isolation system; M, manual method; S, semiautomated method; A, automated method; CY, cell yield [ $\times 10^5$ /ml lipoaspirate]; CV, cell viability [%]; LCY, live cell yield (i.e., number of living cells) [ $\times 10^5$ /ml lipoaspirate]; V<sub>L</sub>, average lipoaspirate volume that was processed [ml]; PT, average processing time [min]; FV, volume of the final cell suspension; --, data not provided or could not be calculated (in case of NLC). (u), Mystem kit (Bi-Medica, Treviolo, Italy); (v), Fatstem (CORIOS Soc. Coop, San Giuliano Milanese, Italy); (w), StromaCell mechanical cell-separation device (MicroAire Aesthetics, Charlottesville, VA, USA); (x), Fastem Corios (CORIOS Soc. Coop, San Giuliano Milanese, Italy).

**Figure 1.**
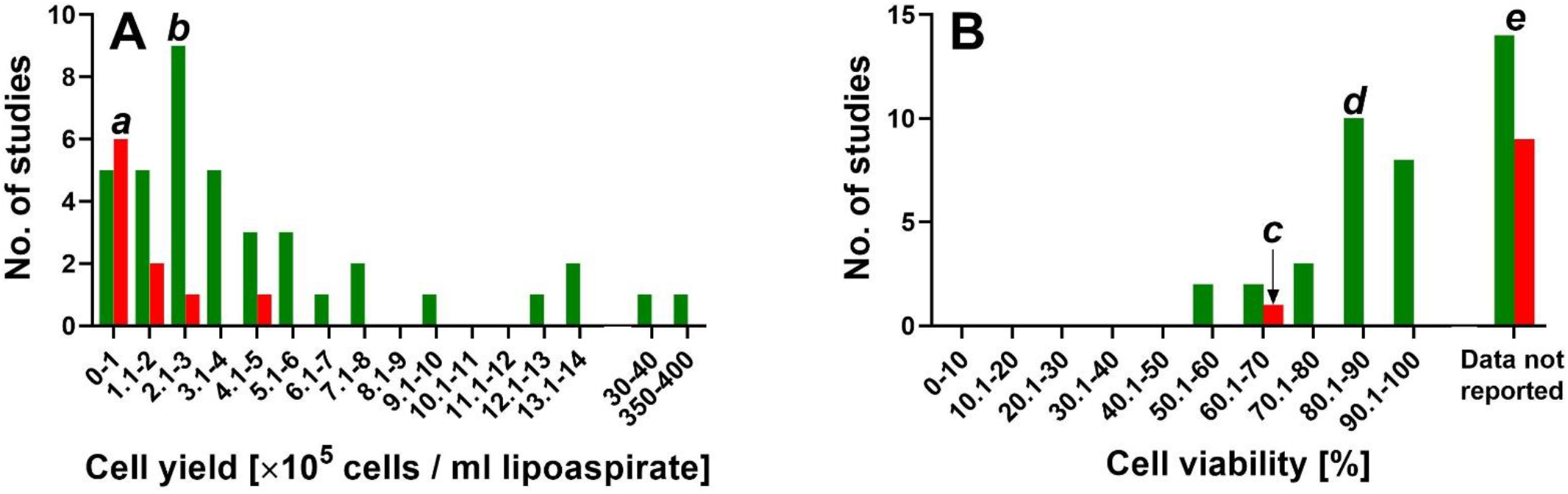
Frequency distributions of cell yield (A) and cell viability (B) data reported in the literature for enzymatic (green) and non-enzymatic (red) methods for isolating ADRCs from adipose tissue. Interpretation of this data is illustrated by the following examples: (i) For six out of ten non-enzymatic methods (60%) cell yield between 0 and 1×10^5^ cells per ml lipoaspirate was reported (“*a*” in Panel A). (ii) For nine out of 39 enzymatic methods (21%) cell yield between 2 ×10^5^ and 3 ×10^5^ cells per ml lipoaspirate was reported (“*b*” in Panel A). (iii) For only one out of ten non-enzymatic methods (10%) cell viability was reported (between 60.1% and 70%) (“*c*” in Panel B). (iv) For ten out of 39 enzymatic methods (20.5%) cell viability between 80.1% and 90% was reported (“*d*” in Panel B). (v) Unfortunately, for nine out of ten non-enzymatic methods (90%) as well as for 14 out of 39 enzymatic methods (36%) no cell viability data were reported (“*e*” in Panel B).

For non-enzymatic methods, cell yield between 0.07×10^5^ [39] and 4.44×10^5^ [57] cells per ml lipoaspirate were reported (Table 2) (mean, 1.1×10^5^; standard deviation, 1.4×10^5^; median, 0.3×10^5^; for 90% of the methods listed in Table 2 cell yield between 0.07×10^5^ and 2.4×10^5^ cells per ml lipoaspirate was reported; Fig 1). Of note, except for one method [53] no cell viability data were reported for non-enzymatic methods (Table 2).

Altogether it follows from these data that (i) both non-enzymatic and enzymatic methods for isolating ADRCs from adipose tissue can result in significant variability in cell yield (Fig 1A); (ii) cell viability data were reported for the majority of enzymatic methods for isolating ADRCs from adipose tissue (25 out of 39 methods; 64%), with cell viability of 80% or higher reported for 18 out of 25 methods (72%) (Fig 1B); (iii) the data reported so far do not allow conclusions regarding cell viability of ADRCs isolated from adipose tissue with non-enzymatic methods (Fig. 1B); and (vi) enzymatic isolation of ADRCs from adipose tissue generally results in higher cell yield than non-enzymatic isolation (Fig. 1A).

The present study tested the following hypotheses: (i) Isolation of ADRCs from human adipose tissue with the Transpose RT system and the proprietary enzymatic Matrase Reagent (both commercially available from InGeneron, Inc., Houston, TX, USA) (thereafter: “Transpose RT / Matrase isolation” and “Transpose RT / Matrase isolated ADRCs”) results in significantly higher cell yield, cell viability and number of living cells per ml lipoaspirate than isolation of ADRCs from human adipose tissue without the use of Matrase Reagent but under otherwise identical processing conditions (thereafter: “Transpose RT / no Matrase isolation” and “Transpose RT/ no Matrase isolated ADRCs”), but does not influence biological characteristics, physiological functions or structural properties of the ADRCs in the final cell suspension. (ii) Transpose RT / Matrase isolation of ADRCs from human adipose tissue results in a higher live cell yield than the methods reported in those studies that are listed in Tables 1 and 2.

## MATERIALS AND METHODS

### Isolation of cells from subcutaneous adipose tissue

Subcutaneous adipose tissue was obtained by a medical practitioner from subjects via lipoaspiration undergoing elective lipoplasty according to standard procedures with informed consent. N=12 subjects with age ranging between 32 and 59 years (Table 3) (and three additional subjects for testing of residual collagenase activity in the final cell suspension) were consented according to the IntegReview IRB approved protocol #200601001 (IntegReview IRB, Austin, TX, USA).

**Table 3.** Subjects’ demographics.

| Subject # | Age | Sex | Race |
| --- | --- | --- | --- |
| 1 | 36 | Female | Caucasian |
| 2 | 59 | Female | Caucasian |
| 3 | 46 | Female | Caucasian |
| 4 | 49 | Female | Caucasian |
| 5 | 37 | Female | Caucasian |
| 6 | 36 | Female | Caucasian |
| 7 | 39 | Female | Black |
| 8 | 36 | Female | Caucasian |
| 9 | 32 | Female | Hispanic |
| 10 | 32 | Female | Hispanic |
| 11 | 38 | Female | Caucasian |
| 12 | 42 | Female | Caucasian |

A sample of recovered lipoaspirate from each subject was divided into two equal parts of 25 ml each, and was processed either with the use of Matrase Reagent (InGeneron) (Transpose RT / Matrase isolation) or only mechanically without the use of Matrase Reagent but under otherwise identical processing conditions (Transpose RT / no Matrase isolation). The Matrase Reagent is a GMP certified, proprietary enzyme blend of collagenase and neutral protease.

Processing was performed as described in the tissue processing procedure section found in the 11011E Transpose RT Instructions for Use (11011-01 IFU; InGeneron, Inc.) (Fig 2), comprising the following steps: (i) The recovered lipoaspirate (25 ml) was loaded together with 2.5 ml reconstituted Matrase (in case of Transpose RT / Matrase isolation) and lactated Ringer solution (preheated to 39° C) into a processing tube up to the MAX FILL line (Fig 2A). (ii) The filled processing tubes were subjected in an inverted position inside the Transpose RT system to repetitive acceleration and deceleration for 30 minutes at 39° C (Fig 2B). (iii) The processed lipoaspirate solution was filtered through a 200 μm filter (Fig 2C) and transferred into a wash tube. After filling the wash tube with saline (room temperature) up to the MAX FILL line, the cells were separated from the rest of the tissue by centrifugation at 600g for 5 minutes at room temperature (Fig 2D). Then, the SVF/ ADRCs (approximately 2 ml) were extracted through a swabable luer vial adapter at the bottom of the wash tube, and the remaining substances (fat, debris and liquid) were discarded (Fig 2E). (v) The cells were returned into the empty wash tube and (after adding fresh saline up to the MAX FILL line) centrifugated again for 5 minutes (Fig 2F). (vi) The previous washing step was repeated (Fig 2G and H). (vii) Finally the concentrated SVF/ ADRCs (approximately 3 ml) were extracted (Fig 2I) and slowly pushed through a luer coupler into a new sterile syringe for further application to the patient.

**Figure 2.**
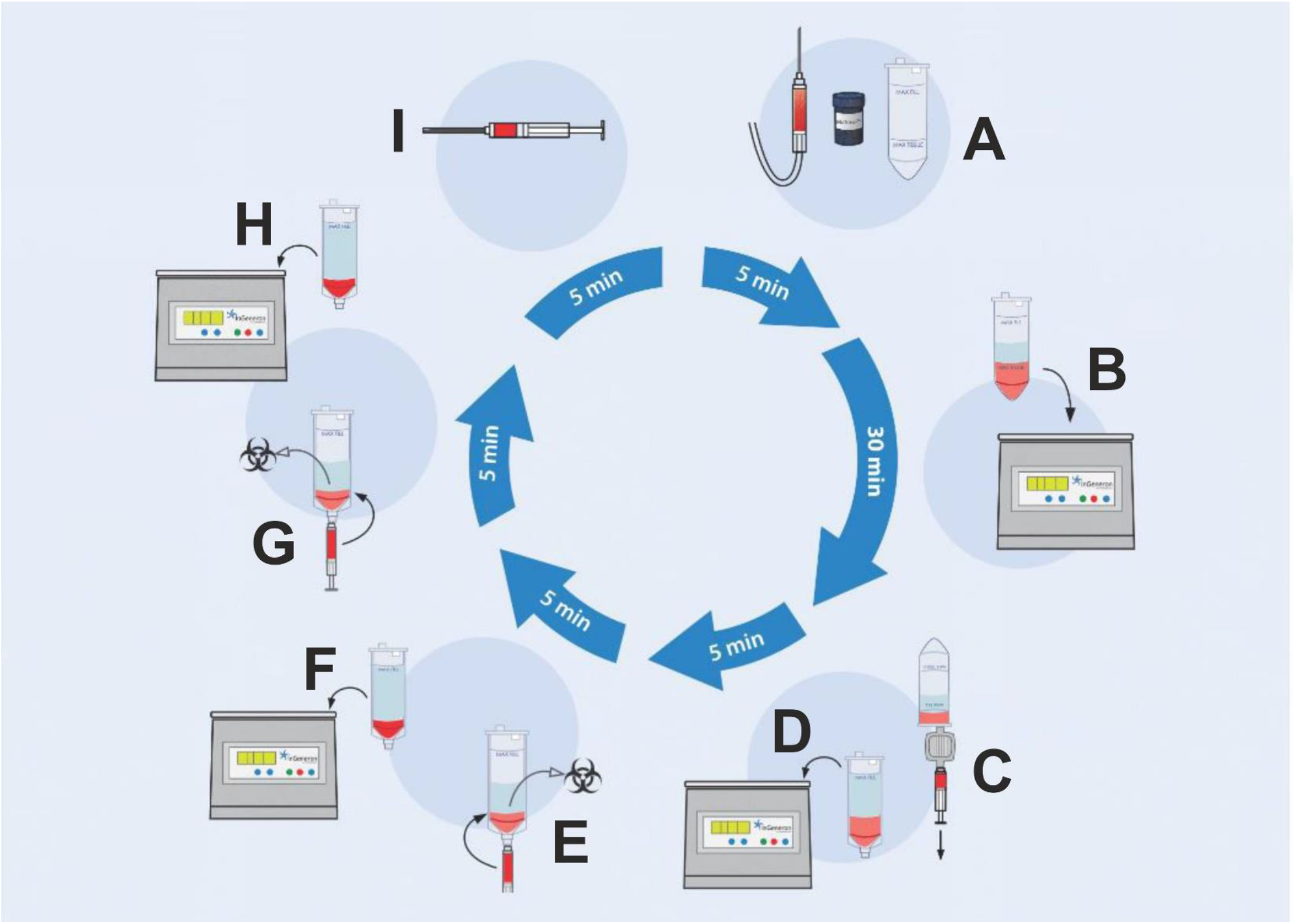
Isolation of ADRCs from human adipose tissue with the Transpose RT system and the Matrase Reagent (both from InGeneron, Inc., Houston, TX, USA). Details are provided in the text.

The final cell suspension was analyzed for cell yield, cell viability, number of living cells per ml lipoaspirate and cell size. The respective differentiation capacity into the three germ layers was assessed for both Transpose RT / Matrase isolated ADRCs and Transpose RT / no Matrase isolated ADRCs.

### SVF yield statistics, expansion, and cryopreservation

Cell counts and viability were determined using the FDA approved NucleoCounter NC-200 device (ChemoMetec Inc., Bohemia, NY, USA) as described by the manufacturer’s protocol.

For expansion, SVF was plated at a cell density of 7×10^6^ in T75 tissue culture flasks in 15 ml of MesenCult MSC basal medium (Stem Cell Technologies, Cambridge, MA, USA) supplemented with MesenCult MSC Stimulatory Supplement, PenStrep (Gibco, Waltham, MA, USA), Fungizone (Life Technologies, Carlsbad, CA, USA) and L-Glutamine (Corning Life Sciences, Tewksbury, MA, USA), at 37° C under 5% CO2. Cells were passed once reaching 75% confluency using 0.25% Trypsin (Sigma-Aldrich, St. Louis, MO, USA) for 5 min at 37° C. Then, cells were plated at a cell density of 7.5×10^5^ in T75 tissue culture flasks in 15 ml of complete MSC medium. Cells were cryopreserved by resuspending ASC cell pellets in Prime-XV MSC FreezIS DMSO-free media (Irving Scientific, Santa Ana, CA, USA) at cell densities between 2×10^6^ and 5×10^6^ cells/ml. The samples were frozen overnight at −80 °C in freezing containers designed to cool at a rate of −1° C/min (Mr. Frosty, Thermo Scientific, Waltham, MA, USA).

### Analysis of colony forming units

The colony forming unit (CFU) assay was performed according to [66]. To this end, freshly isolated SVF/ADRCs from each subject were plated into a 6-well plate (Corning Life Sciences) at two different cell densities. The cells were plated at densities of 50,000 and 100,000 total cells in 2 ml of complete MSC medium in each well; these experiments were repeated in triplicate. Cells were incubated at 37° C under 5% CO2 for 14 days to allow CFUs to form. Medium was changed every 2-4 days. After day 14 the medium was aspirated, the cells were washed twice in PBS and fixed in 2 ml of 10% formalin at room temperature for 30 min with gentle rocking. Cells were then washed three times with DI water and stained with 2 ml hematoxylin (Bio-Rad Laboratories, Hercules, CA, USA) for 15 min at room temperature. The cells were then washed with warm tap water until the wash ran clear. CFUs were quantified by manually counting the entire well; each sample was counted at both cell concentrations and in triplicate. Colonies containing > 50 fibroblast colony-forming units (CFU-F) were counted. CFU-F frequency was calculated by dividing the number of colonies by the number of seeded cells.

### Analysis of mbryoid body formation

Embryoid bodies are defined as spherical clusters of both pluripotent and committed stem cells that can organize in a developmental-specific manner and give rise to mature cells from any differentiation lineage (reviewed in, e.g., [67].

To form embryoid bodies, freshly isolated ADRCs were plated into ultra-low adherent flasks (Corning Life Sciences) at a density of 60,000 cells/cm2 in GMEM (Gibco) supplemented with 2 mM L-Glutamine (Corning Life Sciences), non-essential amino acids (Sigma-Aldrich), B27 (Life Technologies), 0.6% glucose (Sigma-Aldrich), 10 ng/ml human bFGF (Invitrogen Life Technologies), 20 ng/ml human EGF (Life Technologies), 1 U/ml human thrombin (EMD Millipore, Burlington, MA, USA) and 2 μg/ml ciprofloxacin (Sigma-Aldrich). Cells were incubated at 37° C under 5% CO2 and half of the medium was changed every 3 days. The diameter of embryoid bodies was measured on photomicrographs using ImageJ software.

### Differentiation assays

ASCs on their 3rd or 6th passage were assayed for differentiative potential into adipogenic, osteogenic, hepatogenic and neurogenic cell lines.

#### Adipogenic Differentiation

ASCs on their 3^rd^ passage were plated onto a 12 well plate at 40,000 cells per well in 1.5 ml of complete MSC medium and allowed to grow for 2 days. On day 3 all of the medium was aspirated and replaced with either complete MSC medium or StemPro Adipogenic differentiation medium (Life Technologies) and incubated for 2 weeks, changing media every 3-4 days. Then, the presence of intracytoplasmic lipids (triglycerides) was assessed with Oil red-O staining as previously described [60]. Percentage of adipocytes was calculated by microscopic inspection. The percentage of adipocytes was determined by calculating the ratio of Oil red-O positive cells versus total cells.

#### Osteogenic Differentiation

ASCs on their 3^rd^ passage were plated onto a 12 well plate at 20,000 cells per well in 1.5 ml of complete MSC medium and allowed to grow for 2 days. On day 3 all of the medium was aspirated and replaced with either complete MSC medium or StemPro Osteogenic Differentiation medium (Life Technologies) and incubated for 2 weeks, changing media every 3-4 days. Then, the presence of calcific deposits was investigated with Alizarin red staining (Alfa Aesar, Haverhill, MA, USA) as described in the protocol.

#### Hepatogenic Differentiation

ASCs on their 3^rd^ passage were plated onto a 12 well plate at 20,000 cells per well in 1.5 ml of MesenCult MSC medium (Stem Cell Technologies) and allowed to grow for 2 days. Hepatogenic differentiation was achieved using the Human Mesenchymal Stem Cell Hepatogenic Differentiation Medium kit (Cyagen, Santa Clara, CA, USA). Then, the presence of structures containing a high proportion of carbohydrate macromolecules (glycogen, glycoprotein and proteoglycans) was investigated with Periodic Acid Schiff staining (Sigma-Aldrich) as described in the protocol.

#### Neurogenic Differentiation

Nunc Lab-Tek II 4-well chamber slides (Thermo Fisher) were pre-coated with Poly-D-Lysine (Trevigen, Gaithersburg, MD, USA)/Laminin (Gibco) solution (10 μg/ml each) on day 0. Then, 20.000 ASCs on their 6^th^ passage were seeded per well on day 1 and cultured for 24 hours before exposure to Mesencult MSC medium (Stem Cell Technologies) or neurogenic differentiation medium. The latter consisted of Neurobasal Medium (Thermo Fisher) supplemented with 2% B27 (Thermo Fisher) and containing 10 μM Forskolin (Stem Cell Technologies), 5 μg/ml insulin (Sigma-Aldrich), 500 μM 1-methyl-3-isobutylxanthine (IBMX) (Sigma-Aldrich), 50 μM ascorbic acid (Sigma-Aldrich), 10 ng/ml nerve growth factor (NGF) (Thermo Fisher), 1% L-Glutamine (Corning Life Sciences) and 1% Pen/Strep (Gibco). Cells were cultured in neurogenic differentiation medium for 21 days with media changes every 3-4 days.

After 21 days of culturing cells in neurogenic differentiation medium or control medium, they were fixed with 10% formalin (Sigma-Aldrich). Then, cells were treated with PBS containing 0.3% Triton X-100 (Sigma-Aldrich) and 5% normal goat serum (Jackson Immunoresearch, West Grove, PA, USA) at room temperature for 1 h to block non-specific binding sites prior to the addition of primary antibodies. Afterwards, cells were incubated with primary antibodies against microtubule-associated protein 2 (MAP2) (rabbit polyclonal, 1:100, ab32454; Abcam, Cambridge, UK) or Beta III Tubulin (β3TUB) (rabbit monoclonal, 1:200, β3-Tubulin (D71G9) XP Rabbit mAb #5568; Cell Signaling Technology, Danvers, MA, USA) for 2 h at room temperature. (MAP2 is a neuron-specific cytoskeletal protein that is used as a marker of neuronal phenotype [68], and β3TUB is a tubulin thought to be specifically involved during differentiation of neuronal cell types [69].) Antibodies were diluted in antibody dilution buffer (PBS containing 1% bovine serum albumin (VWR, Radnor, PA, USA) and 0.3% Triton X-100 (Sigma-Aldrich)). After incubation with primary antibodies, the cells were washed three times with PBS, followed by incubation with goat anti-rabbit secondary antibody conjugated with Alexa Fluor 594 (Thermo Fisher) for 1 h at room temperature. Then, cells were again washed three times with PBS, followed by counterstaining with DAPI (Sigma-Aldrich). Finally, chambers were removed from the slides, and coverslips were mounted with Vectashield Antifade Mounting Medium (Vector Laboratories, Burlingame, CA, USA).

PC-12 cells (a model system for neuronal differentiation [70]) were used as positive controls. PC-12 cells were seeded onto COL IV (Santa Cruz, Santa Cruz, CA, USA) coated Nunc Lab-Tek II 4-well chamber slides, with 8,000 cells per well in neurogenic differentiation medium. Cells were cultured for 6 days and fixed with 10% formalin. Immunofluorescent detection of MAP2 and β3TUB was performed as described for ASCs.

### RNA isolation and Quantitative PCR

RNA isolation was performed using Trizol (Life Technologies) in accordance with the manufacturer’s protocol. Total RNA was purified using the Direct-Zol RNA miniprep kit (Zymo Research, Irvine, CA, USA) as described in the protocol. cDNA was generated using iScript Reverse Transcriptase SuperMix (Bio-Rad Laboratories). Then, relative mRNA levels of Oct4 (a transcription factor associated with self-renewal [72], Klf4 (a marker of stemness [72]) and Hes1 (another known stem cell marker [73]) were measured using the SsoAdvanced Universal SYBR Green Supermix (Bio-Rad Laboratories) according to the manufacturer’s protocols in a Bio-Rad CFX96 Real-Time PCR detection system (Bio-Rad Laboratories). All samples were analyzed in triplicate. Qualification of RNA and cDNA was performed using a NanoDrop spectrometer (Thermo Fisher Scientific). Primer probe sets were custom oligos (Sigma-Adrich) (Table 4).

**Table 4.**
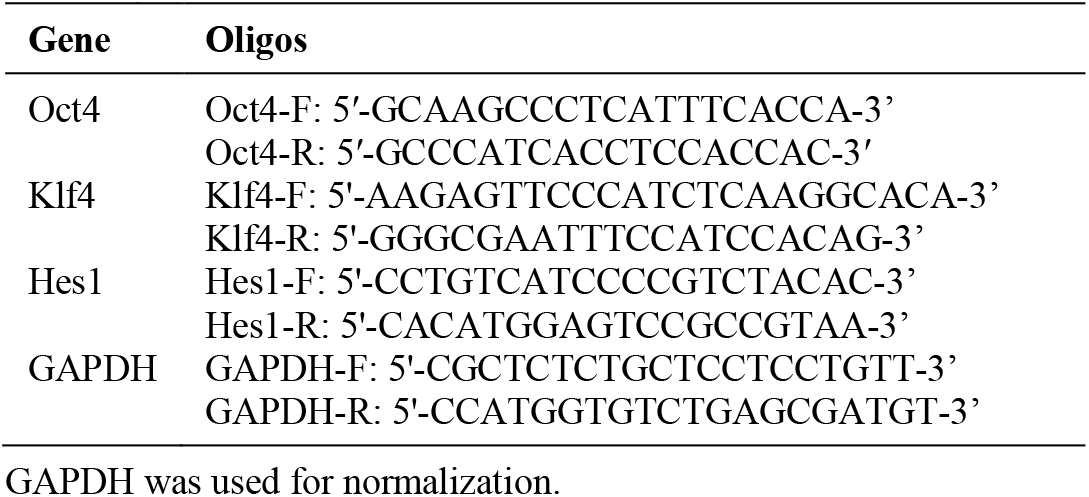
Primer probe sets used in the present study.

Amplification curves and melting curves were illustrated by CFX Maestro software (Version 1.1; Bio-Rad Laboratories). Relative mRNA levels were determined by the ΔΔCt quantification method using the CFX Maestro software. Baseline correction and threshold setting were performed using the automatic calculation offered by the software.

### Testing for residual collagenase activity in cell preparations prepared with the use of Matrase Reagent

In this experiment ADRCs were isolated from lipoaspirate from n=3 additional subjects using the ARC system (InGeneron; a slight modification of the Transpose RT system, adapted for additional use with larger portions of adipose tissue) and Matrase Reagent following the manufacturer’s instructions for use. The resulting cell preparations were tested for collagenase activity using a commercially available assay (EnzChek Gelatinase/Collagenase Assay Kit; Invitrogen, Carlsbad, CA, USA) following the manufacturer’s instructions for use.

### Microscopy

Images were obtained using an Axio Vert.A1 microscope (Carl Zeiss Microscopy, Jena, Germany) equipped with LD A-Plan 10x/0,25 and LD A-Plan 20x/0.30 objectives (Carl Zeiss Microscopy), Leica DMC4500 camera (2560 × 1920 pixels; Leica Microsystems, Wetzlar, Germany) and Leica Application Suite software (version X 3.3.3.16958; Leica). The final figures were constructed using Corel Photo-Paint X7 and Corel Draw X7 (both versions 17.5.0.907; Corel, Ottawa, Canada). No adjustments of contrast and brightness were made.

### Statistical analysis

Mean and standard error of the mean (SEM) were calculated for all variables. The D’Agostino and Pearson omnibus normality test was used to determine whether the distribution of the investigated variables of the Transpose RT / Matrase isolated cells and the Transpose RT / no Matrase isolated cells were consistent with a Gaussian distribution. Differences between the groups of cells were tested with nonparametric Wilcoxon matched-pairs signed rank test. In all analyses, an effect was considered statistically significant if its associated p value was smaller than 0.05. Calculations were performed with GraphPad Prism (Version 5; Graph Pad Software, San Diego, CA, USA).

## RESULTS

### Cell yield, cell viability, number of living cells per ml lipoaspirate and cell size

Compared to Transpose RT / no Matrase isolation, Transpose RT / Matrase isolation of ADRCs from lipoaspirate resulted in the following, statistically significant differences in the final cell suspension (all values given as mean ± SEM): (i) approximately nine times higher cell yield (7.2×10^5^ ± 0.90×10^5^ Transpose RT / Matrase isolated ADRCs per ml lipoaspirate vs. 0.84×10^5^ ± 0.10×10^5^ Transpose RT / no Matrase isolated ADRCs per ml lipoaspirate; p < 0.001; n=12 matched pairs of samples) (Fig 3A); (ii) approximately 41% higher mean cell viability (85.9% ± 1.1% in case of Transpose RT / Matrase isolated ADRCs vs. 61.7% ± 2.6% in case of Transpose RT / no Matrase isolated ADRCs; p < 0.001; n=12 matched pairs of samples) (Fig 3B); and (iii) approximately twelve times higher mean number of living cells per ml lipoaspirate (6.25×10^5^ ± 0.79×10^5^ Transpose RT / Matrase isolated ADRCs per ml lipoaspirate vs. 0.52×10^5^ ± 0.08×10^5^ Transpose RT / no Matrase isolated ADRCs per ml lipoaspirate; p < 0.001; n=12 matched pairs of samples each) (Fig 3C).

**Figure 3.**
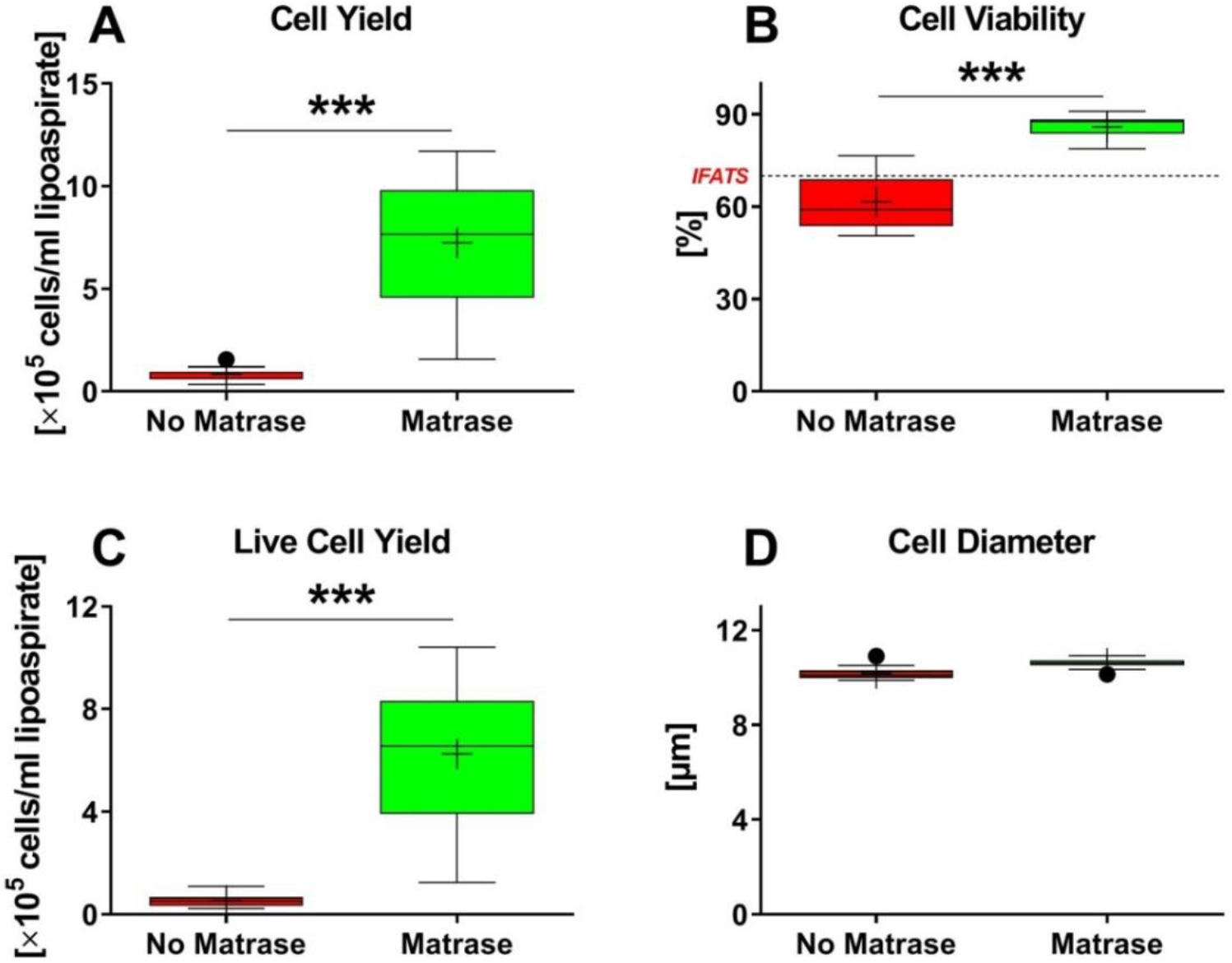
Results of quantitative analysis of Transpose RT / no Matrase isolated ADRCs and Transpose RT / Matrase isolated ADRCs. The panels show Tukey boxplots of the number of cells isolated per ml lipoaspirate (**A**), the relative number of viable cells (**B**), the number of viable cells isolated per ml lipoaspirate (**C**) and the diameter of cells (**D**) of Transpose RT / no Matrase isolated ADRCs (red bars) and Transpose RT / Matrase isolated ADRCs (green bars). In (**B**) the threshold of 70% viable cells established by the International Federation for Adipose Therapeutics and Science (IFATS) [71] is indicated by a dashed line. Results of Wilcoxon matched-pairs signed rank test are indicated (n=12 paired samples each). ***, p < 0.001.

Of importance, the mean relative number of viable cells obtained by Transpose RT / Matrase isolation (85.9%) exceeded the proposed minimum threshold for the viability of cells in the SVF of 70 % established by the International Federation for Adipose Therapeutics and Science (IFATS) [71], whereas the mean relative number of viable cells obtained by Transpose RT / no Matrase isolation (61.7%) did not (Fig 3B).

The difference in mean cell diameter between Transpose RT / no Matrase isolated ADRCs (10.2μm ± 0.1μm) and Transpose RT / Matrase isolated ADRCs (10.6μm ± 0.1μm) was only approximately 4% and did not reach statistical significance (p = 0.05; n=12 matched pairs of samples) (Fig 3D).

Accordingly, both the number and viability of cells in the final cell suspension were statistically significantly higher after Transpose RT / Matrase isolation of ADRCs from human adipose tissue than after Transpose RT / no Matrase isolation.

### Colony-Forming Unit assay

ASCs derived from Transpose RT / Matrase isolated ADRCs formed on average 16 times more CFUs per ml lipoaspirate (4973±836; mean ± SEM) than ASCs derived from Transpose RT / no Matrase isolated ADRCs (307±68) (p = 0.002; n=10 matched pairs of samples) (Fig 4).

**Figure 4.**
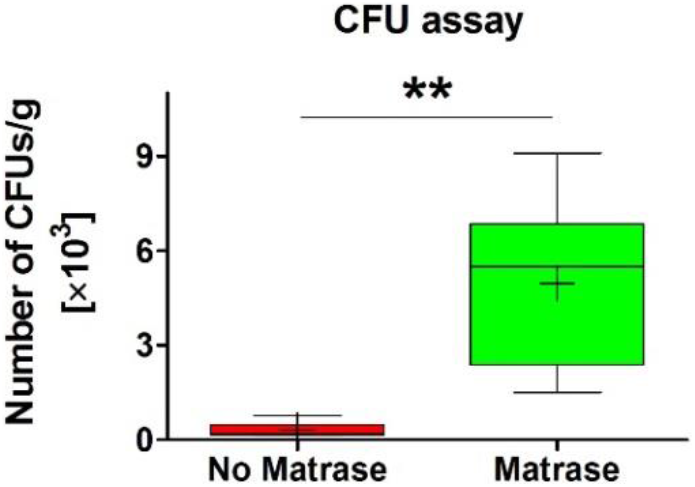
Results of colony forming unit assay. The panel shows Tukey boxplots of the number of colony forming units per ml lipoaspirate formed by ASCs derived from Transpose RT / no Matrase isolated ADRCs (red bars) and from Transpose RT / Matrase isolated ADRCs (green bars) after culturing for 14 days in complete MSC media. Results of Wilcoxon matched-pairs signed rank test are indicated (n=10 paired samples). **, p < 0.005.

### Expression of regenerative cell-associated genes in adipose-derived stem cells derived from uncultured, autologous adipose-derived regenerative cells

Embryoid body formation was observed after culturing ASCs for seven days in serum free media (Fig 5). The embryoid bodies had a spherical appearance and defined borders. The majority of the embryoid bodies were small in diameter (<100 μm), some were medium (100-200 μm), and a few were large (>200 μm) in diameter. No statistically significant differences were observed in the formation or the size of the embryoid bodies derived from Transpose RT / no Matrase isolated ADRCs and from Transpose RT / Matrase isolated A (Fig 6).

**Figure 5.**
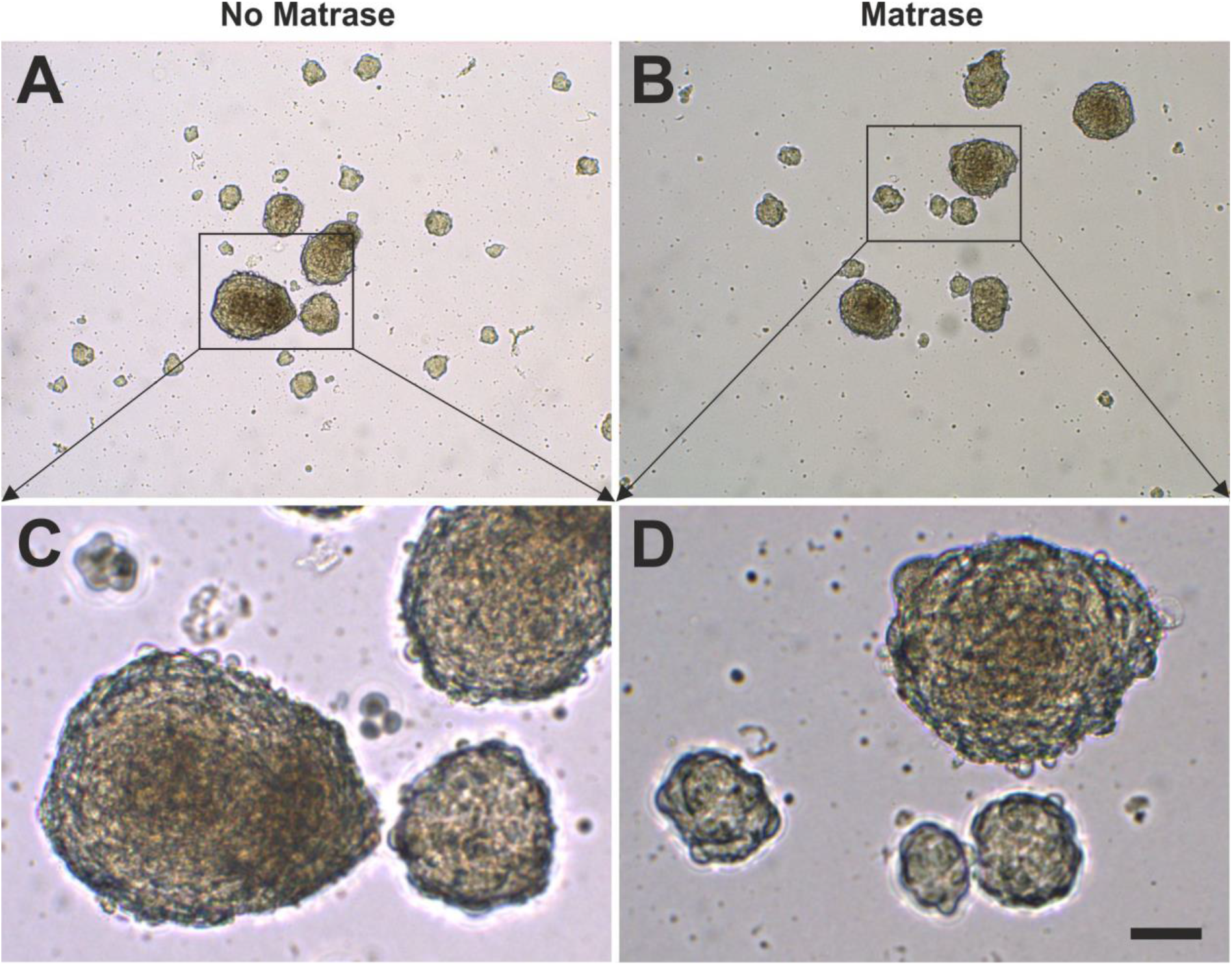
Formation of embryoid bodies. The panels show embryoid bodies that were formed after culturing ASCs derived from Transpose RT / no Matrase isolated ADRCs (**A**, **C**) and from Transpose RT / Matrase isolated ADRCs (**B**, **D**) for seven days in serum-free media. The scale bar in (D) represents 100 μm in (A, B) and 50 μm in (B, D).

**Figure 6.**
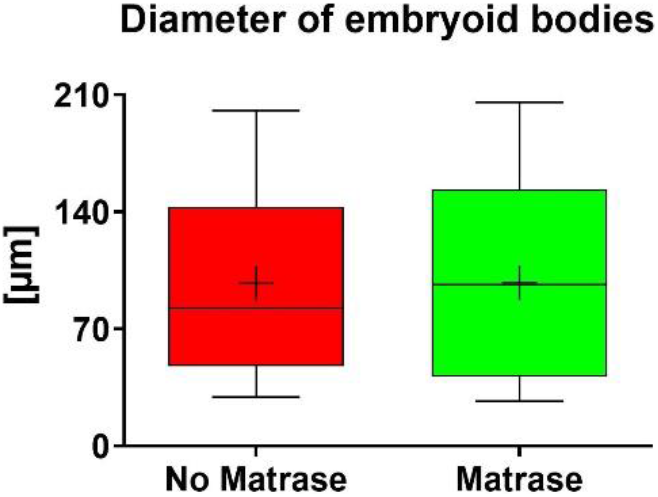
Quantitative analysis of the size of embryoid bodies. The panel shows Tukey boxplots of the diameter of embryoid bodies that were formed after culturing ASCs derived from Transpose RT / no Matrase isolated ADRCs (red bars) and from Transpose RT / Matrase isolated ADRCs (green bars) for seven days in serum-free media. The Wilcoxon matched-pairs signed rank test showed no statistically significant differences between the groups (p = 0.109; n=4 paired samples).

Furthermore, it was found that the use of Matrase Reagent in the process of isolating cells from lipoaspirate had no statistically significant impact on the relative levels of mRNA for the regenerative cell-associated genes Oct-4, Klf4 and Hes1 for both conventional monolayer cultures (Fig 7A,C,E) and embryoid body cultures (Fig 7B,D,F) (mean and SEM of relative gene expression values as well as corresponding p-values are summarized in Table 5).

**Figure 7.**
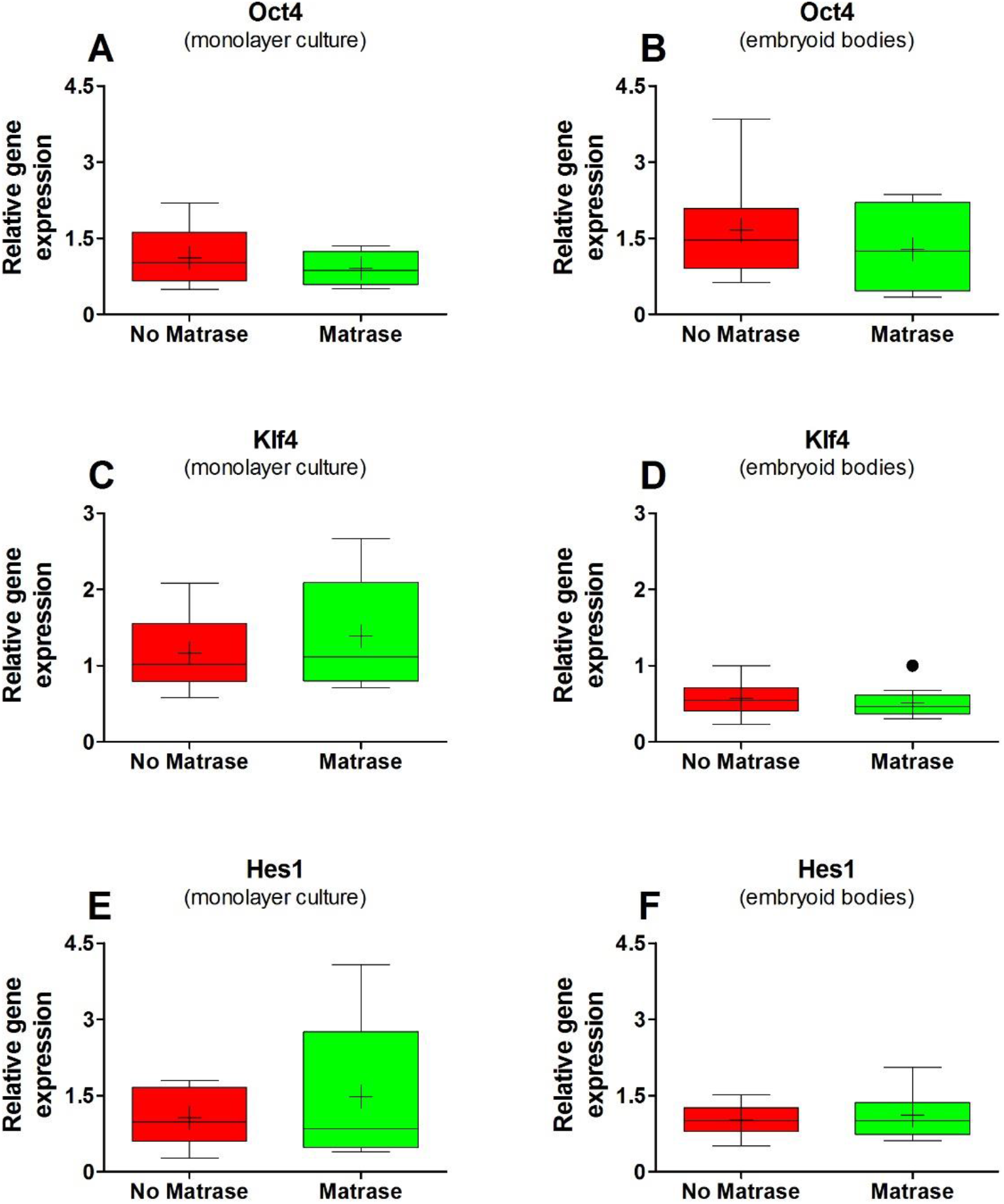
Results of gene expression analysis. The panels show Tukey boxplots of relative gene expression (arbitrary units) of Oct4 (**A**, **B**), Klf4 (**C**, **D**) and Hes3 (**E**, **F**) of ASCs in conventional monolayer cultures (A, C, E) or obtained from embryoid bodies (B, D, F) after culturing Transpose RT / no Matrase isolated ADRCs (red bars) or Transpose RT / Matrase isolated ADRCs (green bars), respectively. The Wilcoxon matched-pairs signed rank test showed no statistically significant differences between the groups (p > 0.05; n=8 paired samples each).

**Table 5.**
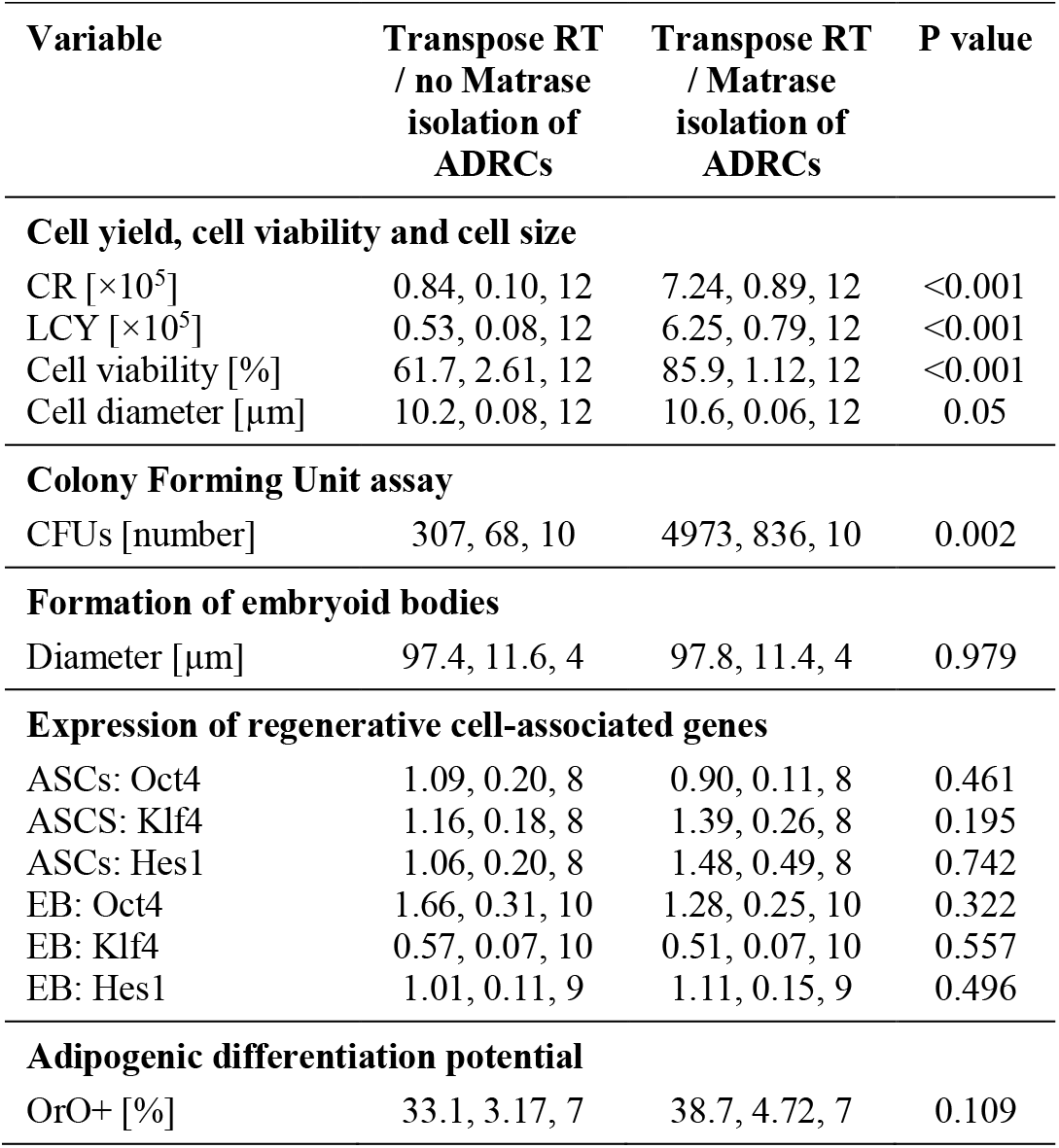
Results of statistical analysis. The table shows the results of all statistical analyses that were performed in this study. All data are provided as {mean, standard error of the mean [SEM], number of paired samples}. P-values were obtained using nonparametric Wilcoxon matched-pairs signed rank test. Cell recovery and live cell yield data represent numbers of cells per gram processed lipoaspirate; relative gene expression data are provided in arbitrary units. Calculations were performed with GraphPad Prism (Version 5; Graph Pad Software, San Diego, CA, USA). Note: the different numbers of paired samples in the gene expression studies are related to the quantity and amount of mRNA generated for each sample. Samples that yielded either insufficient amount of mRNA or lower quality mRNA were not analyzed for the relative level of gene expression. Abbreviations: CR, cell recovery; LCY, live cell yield; CFUs, colony forming units; ASCs, adipose-derived stem cells; EB, embryoid bodies; OrO+, No. of Oil red-O positive cells

Accordingly, the use of Matrase Reagent in processing lipoaspirate with the InGeneron Transpose RT System did not alter expression of regenerative cell-associated genes in the final cell suspension.

### Differentiation potential of adipose-derived stem cells

#### Adipogenic differentiation potential

ASCs on their 3^rd^ passage (derived from both Transpose RT / Matrase isolated ADRCs and Transpose RT / no Matrase isolated ADRCs) were cultured for two weeks in adipogenic differentiation medium or control medium. Then, the presence of intracytoplasmic lipids (triglycerides) was assessed with Oil red-O staining, and relative numbers of Oil red-O positive cells were evaluated. It was found that the use of Matrase Reagent in the process of isolating ADRCs from lipoaspirate had no impact on the visual appearance of the cells after induction of adipogenic differentiation (Fig 8), and no statistically significant impact on the relative number of Oil red-O positive cells (Fig 9).

**Figure 8.**
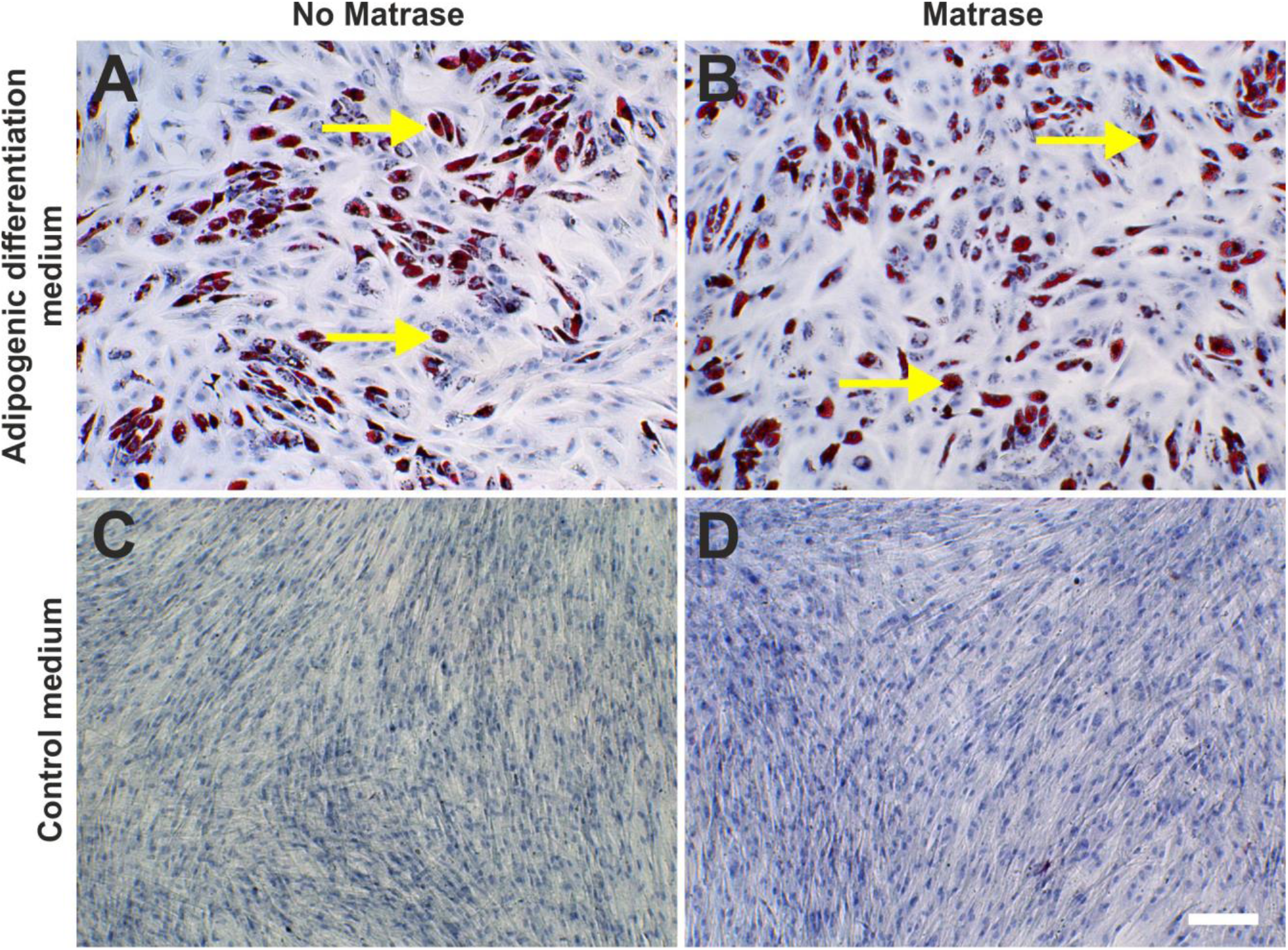
Adipogenic differentiation potential of ADRCs. The panels show the results of culturing ASCs on their 3^rd^ passage (derived from Transpose RT / no Matrase isolated ADRCs (**A**, **C**) or Transpose RT / Matrase isolated ADRCs (**B**, **D**), respectively) for two weeks in adipogenic differentiation medium (A, B) or control medium (C, D). The presence of intracytoplasmic lipids (triglycerides) was assessed with Oil red-O staining; cells were counterstained with hematoxylin. The yellow arrows indicate single Oil red-O positive cells. The scale bar in (D) represents 100 μm in (A-D).

**Figure 9.**
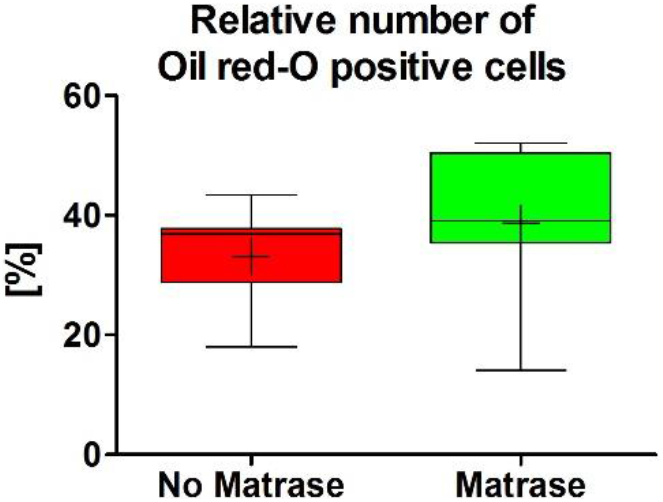
Quantitative analysis of adipogenic differentiation potential of ADRCs. The panel shows Tukey boxplots of the relative number of Oil red-O positive cells obtained after culturing ASCs on their 3rd passage (derived from Transpose RT / no Matrase isolated ADRCs (red bars) or Transpose RT / Matrase isolated ADRCs (green bars), respectively) for two weeks in adipogenic differentiation medium. The Wilcoxon matched-pairs signed rank test showed no statistically significant differences between the groups (p = 0.109; n=7 paired samples).

#### Osteogenic differentiation potential

ASCs on their 3^rd^ passage (derived from both Transpose RT / Matrase isolated ADRCs and Transpose RT / no Matrase isolated ADRCs) were cultured for two weeks in osteogenic differentiation medium or control medium. Then, the presence of calcific deposits was investigated with Alizarin red staining. It was found that the use of Matrase Reagent in the process of isolating ADRCs from lipoaspirate had no impact on the visual appearance of the cells after induction of osteogenic differentiation (Fig 10).

**Figure 10.**
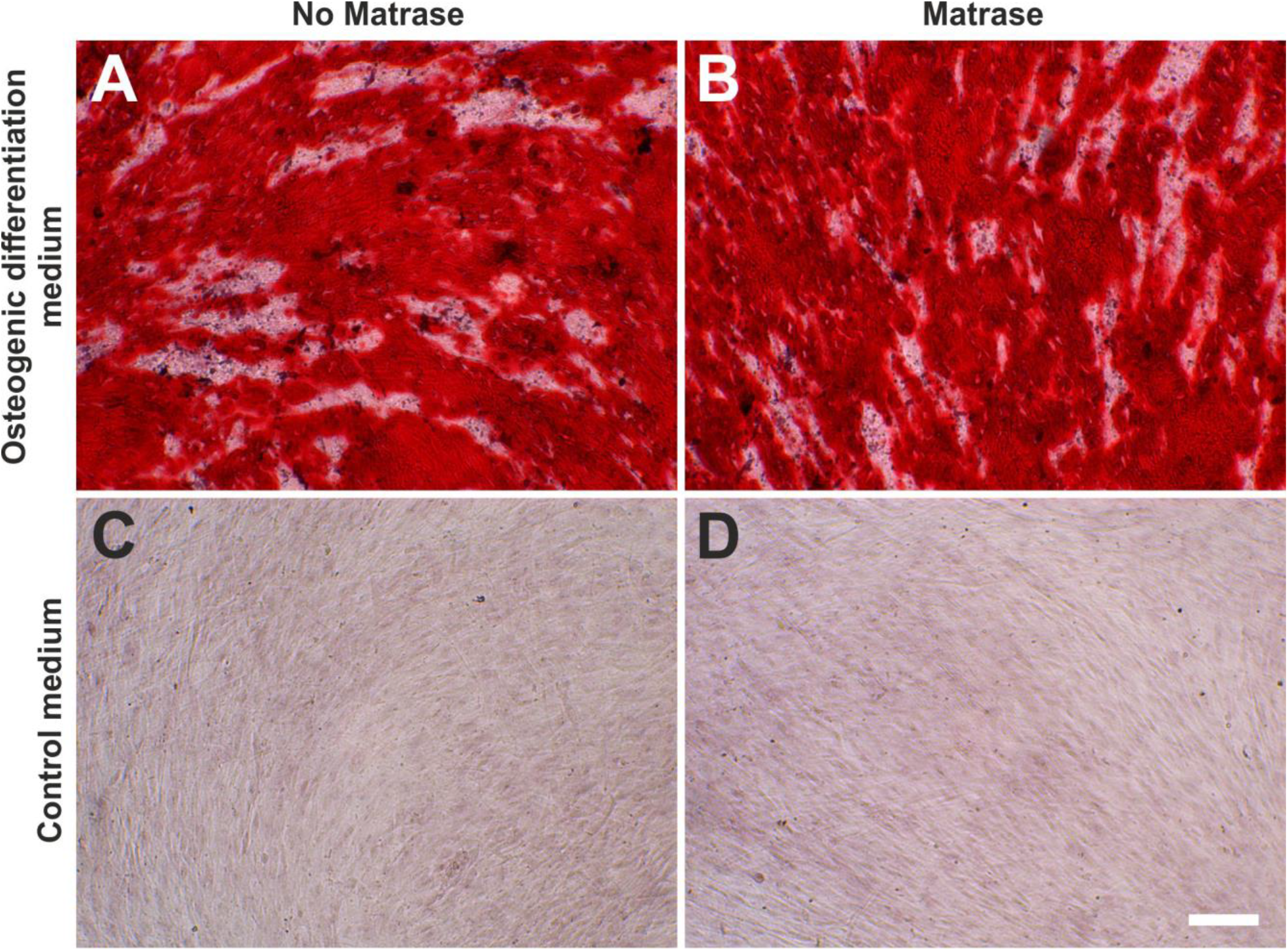
Osteogenic differentiation potential of ADRCs. The panels show the results of culturing ASCs on their 3^rd^ passage (derived from Transpose RT / no Matrase isolated ADRCs (**A**, **C**) or Transpose RT Matrase isolated ADRCs (**B**, **D**), respectively) for two weeks in osteogenic differentiation medium (A, B) or control medium (C, D). The presence of calcific deposits was investigated with Alizarin red staining; cells were counterstained with hematoxylin. Cells of an osteogenic lineage are stained bright to deep red and easily visible as dense red patches. The scale bar in (D) represents 100 μm in (A-D).

#### Hepatogenic differentiation potential

ASCs on their 3^rd^ passage (derived from both Transpose RT / Matrase isolated ADRCs and Transpose RT / no Matrase isolated ADRCs) were cultured for ten days in hepatogenic differentiation medium or control medium. Then, the presence of structures containing a high proportion of carbohydrate macromolecules (glycogen, glycoprotein and proteoglycans) was investigated with Periodic Acid Schiff staining. It was found that the use of Matrase Reagent in the process of isolating ADRCs from lipoaspirate had no impact on the visual appearance of the cells after induction of hepatogenic differentiation (Fig 11.

**Figure 11.**
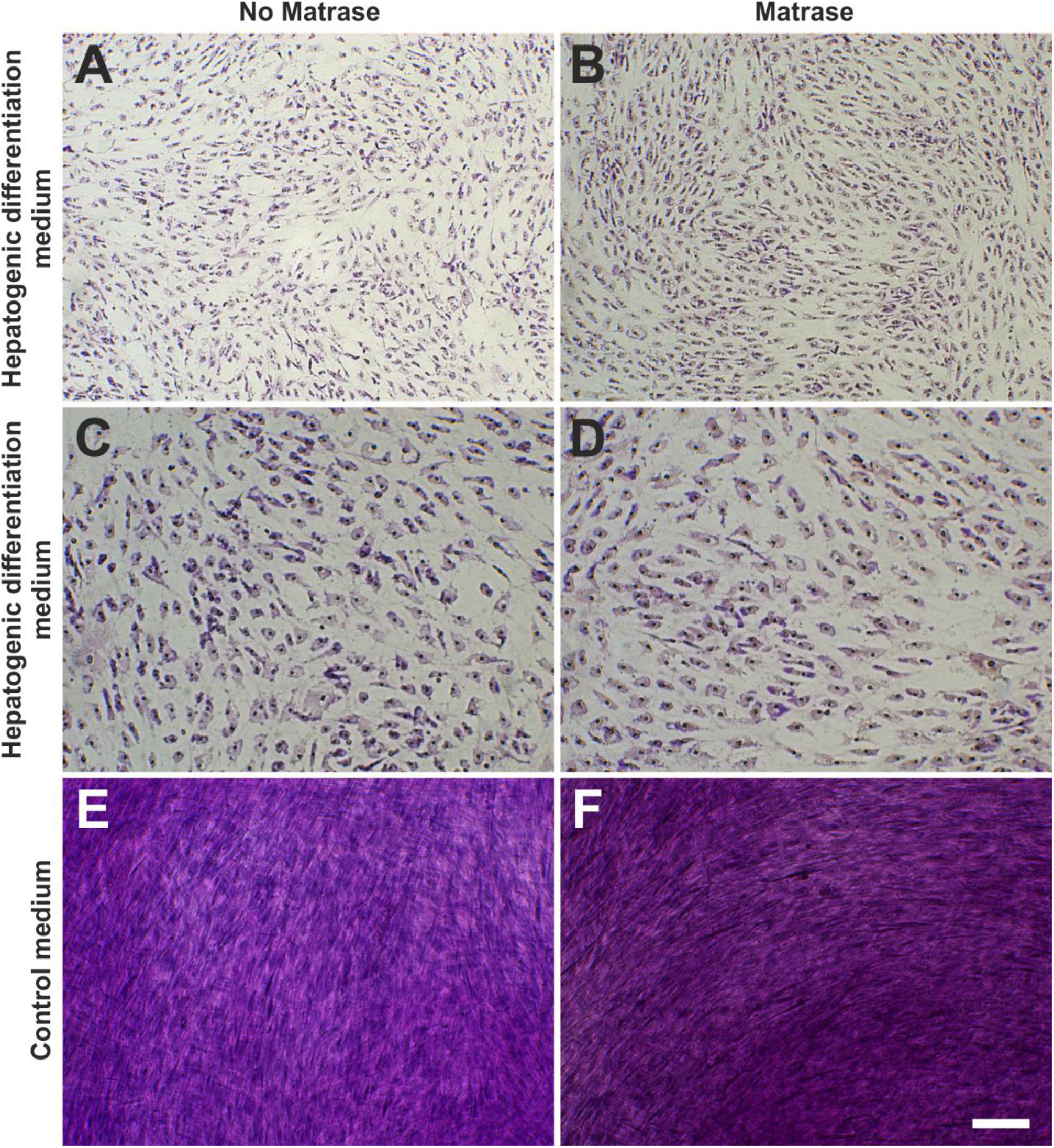
Hepatogenic differentiation potential of ADRCs. The panels show the results of culturing ASCs on their 3^rd^ passage (derived from Transpose RT / no Matrase isolated ADRCs (**A**, **C**, **E**) or Transpose RT / Matrase isolated ADRCs (**B**, **D**, **F**), respectively) for ten days in hepatogenic differentiation medium (A-D) or control medium (E, F). The presence of structures containing a high proportion of carbohydrate macromolecules (glycogen, glycoprotein and proteoglycans) was investigated with Periodic Acid Schiff staining; cells were counterstained with hematoxylin. As a result of induction of hepatogenesis the morphology of the cells changed from a fibroblastic spindle shape to a rather polygonal shape typically associated with hepatocytes. The scale bar in (F) represents 100 μm in (A, B, E, F) and 50 μm in (C, D).

#### Neurogenic differentiation potential

ASCs on their ^6th^ passage (derived from both Transpose RT / Matrase isolated ADRCs and Transpose RT / no Matrase isolated ADRCs) were cultured for three week in neurogenic differentiation medium or control medium. Then, the morphology of the cells was investigated with phase contrast microscopy, and expression of MAP2 and β3TUB with immunofluorescence. It was found that as a result of induction of neurogenesis, the cells showed key characteristics of a neuronal phenotype, i.e., slender processes (Fig 12) and expression of both MAP2 and β3TUB (Fig 13). PC12 cells served as positive controls (Fig 14). The use of Matrase Reagent in the process of isolating ADRCs from lipoaspirate had no impact on the visual appearance of the cells after induction of neurogenic differentiation (Figs 12 and 13).

**Figure 12.**
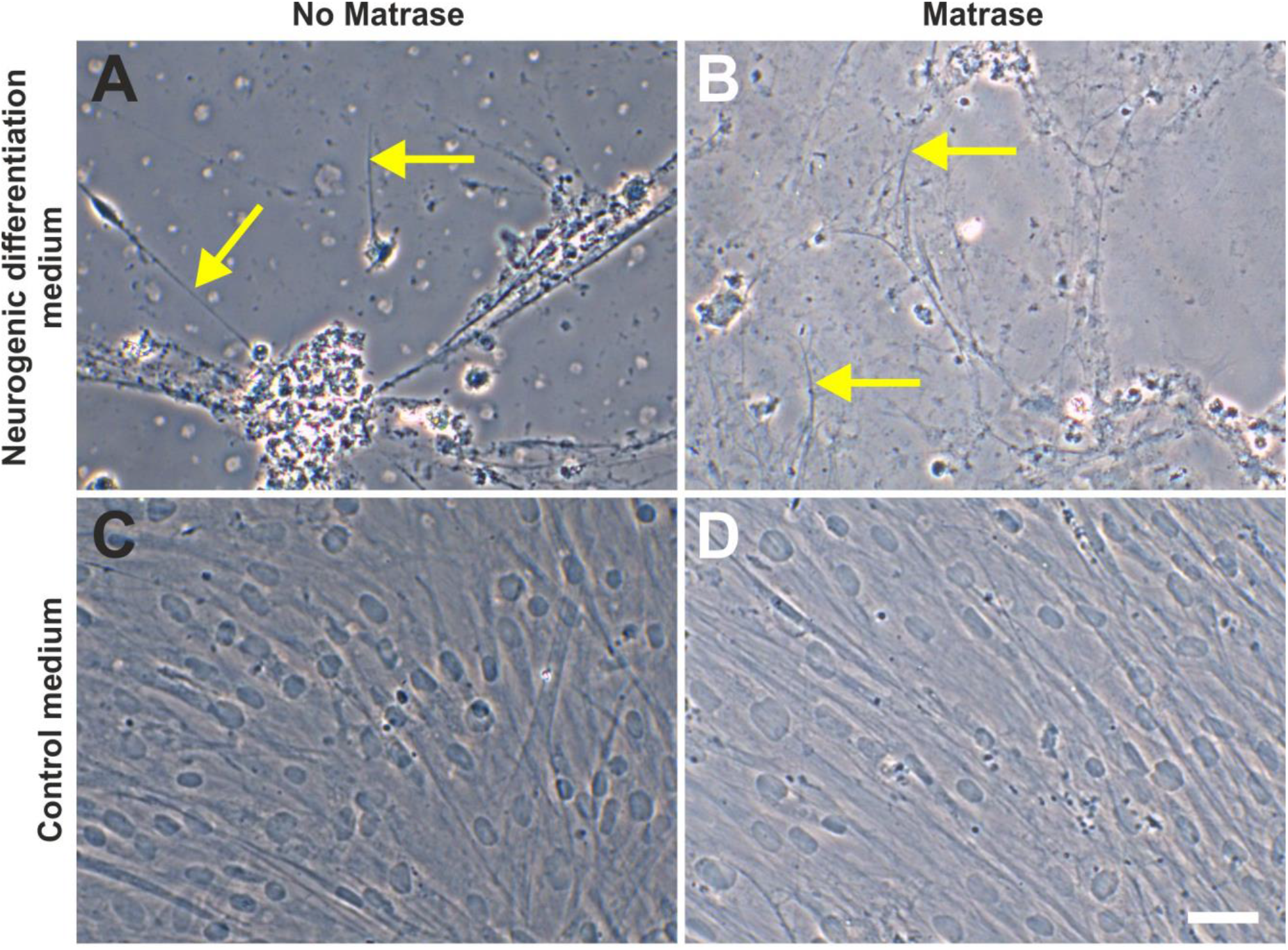
Neurogenic differentiation potential of ADRCs (1). The panels show the results of culturing ASCs on their 6^th^ passage (derived from Transpose RT / no Matrase isolated ADRCs (**A**, **C**) or Transpose RT / Matrase isolated ADRCs (**B**, **D**), respectively) for three weeks in neurogenic differentiation medium (A, B) or control medium (C, D). Cells were imaged with phase contrast microscopy. As a result of induction of neurogenesis, the cells developed characteristic, slender processes (arrows in A, B). The scale bar in (D) represents 50 μm in (A-D).

**Figure 13.**
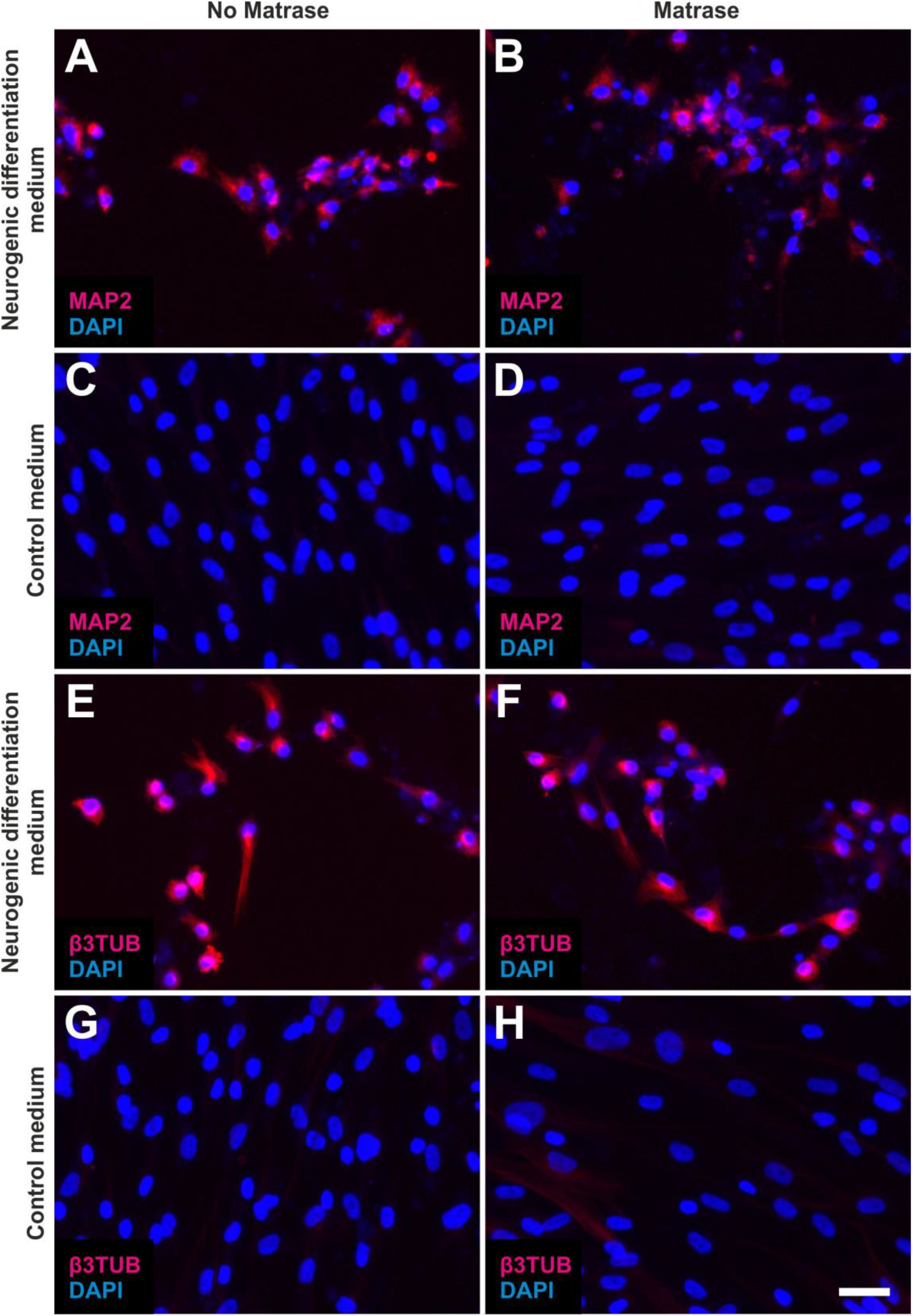
Neurogenic differentiation potential of ADRCs (2). The panels show the results of culturing ASCs on their 6^th^ passage (derived from Transpose RT / no Matrase isolated ADRCs (A, C, E, G) or Transpose RT / Matrase isolated ADRCs (B, D, F, H), respectively) for three weeks in neurogenic differentiation medium (A, B, E, F) or control medium (C, D, G, H). Cells were processed with immunofluorescence for the detection of respectively microtubule-associated protein 2 (MAP2) (A-D) or beta III Tubulin (β3TUB) (E-H), and were counterstained with DAPI. As a result of induction of neurogenesis, the cells expressed both MAP2 (A, B) and β3TUB (E, F). The scale bar in (H) represents 50 μm in (A-H).

**Figure 14.**
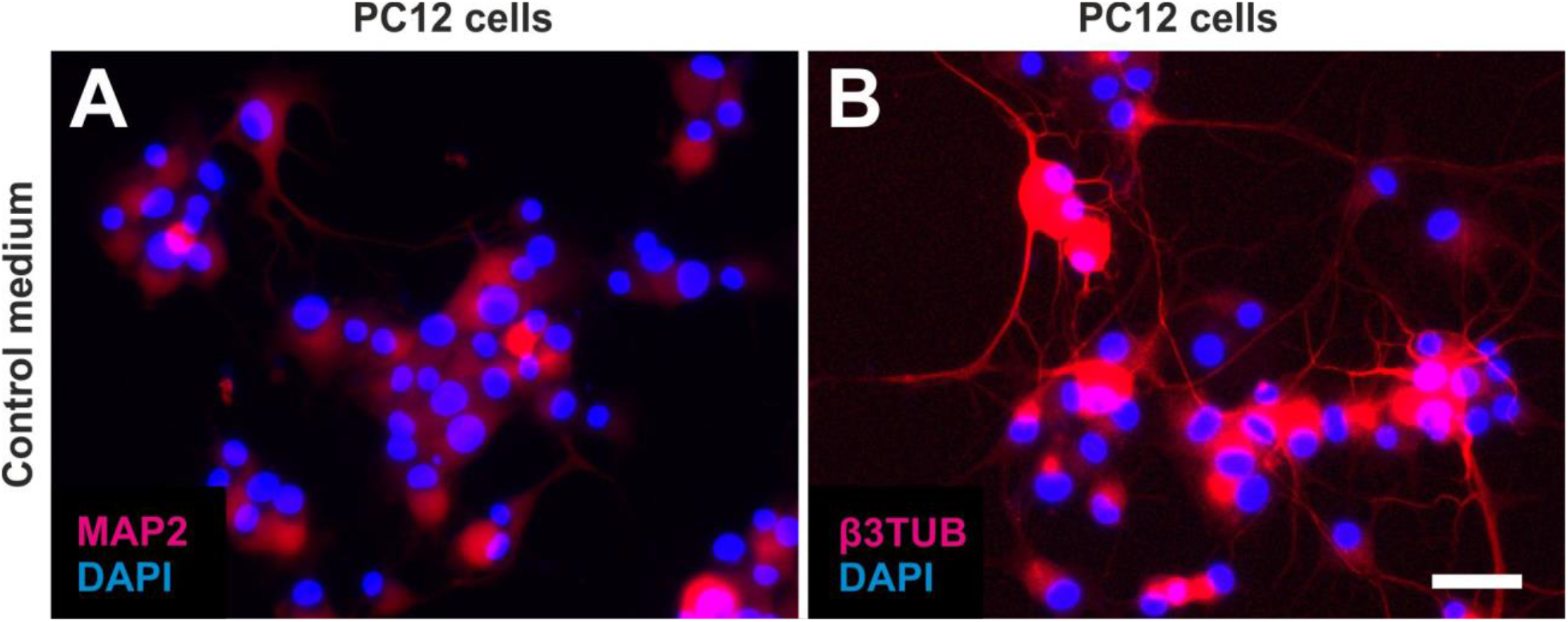
Expression of microtubule-associated protein 2 (MAP2) and beta III tubulin (β3TUB) by PC12 cells as positive control. The panels show the results of culturing PC12 cells for 6 days in neurogenic differentiation medium. Cells were processed with immunofluorescence for the detection of respectively MAP2 (A) or β3TUB (B), and were counterstained with DAPI. The scale bar in (B) represents 50 μm in (A, B).

### Residual collagenase activity in cell preparations prepared with the use of Matrase Reagent

It was found that the collagenase activity was below the detection limit of the used assay (Fig 15).

**Figure 15.**
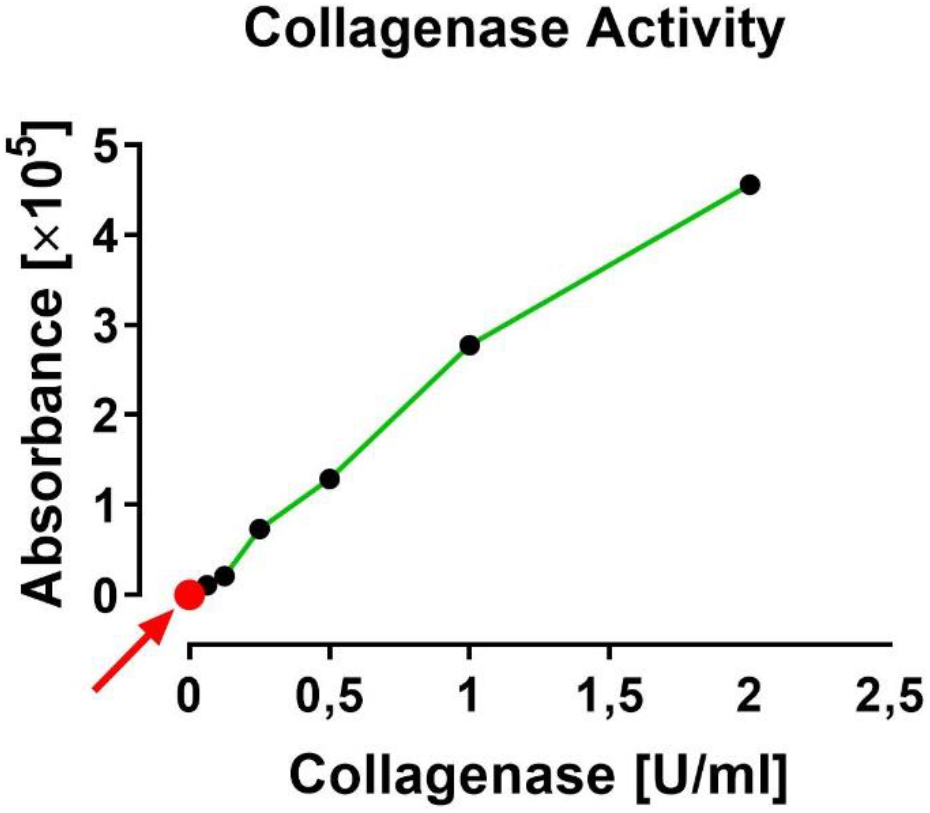
Measurement of collagenase activity in cell preparations that were prepared with the use of Matrase Reagent following the manufacturer’s instructions for use. No collagenase activity was detected in cell preparations (the red dot at Y = X = 0 [indicated by the red arrow] represents the mean of two independent measurements for each sample). The standard curve for the used assay (EnzChek Gelatinase/Collagenase Assay Kit E12055; Invitrogen) is depicted by the green line. Because this high-sensitivity assay can detect enzyme activity down to a minimum concentration of 2×10^−3^ U/ml, activity of the enzyme in the final cell preparation was not present or less than two thousands of a unit per ml.

## DISCUSSION

Aronowitz and colleagues [43] proposed to judge a system or method for isolating ADRCs from adipose tissue by the following factors: nucleated cell count, nucleated cells per milliliter of tissue processed, cellular viability, level of residual enzymatic activity, data from flow cytometry and CFU-F assay, infection control, ease of use, cost to operate and processing time.

There are currently several systems offered and methods described in the literature for clinical therapeutic usage (or that are under clinical evaluation) that process adipose tissue with (Table 1) or without (Table 2) the use of enzymes. It was the aim of this study to evaluate the effect on cells, efficiency and safety regarding cell viability in direct comparison between processing human lipoaspirate with or without the use of an enzyme under otherwise identical processing conditions. InGeneron’s Transpose RT system utilizes a proprietary method for isolating ADRCs from adipose tissue. Specifically, this method uses the enzymatic activity of Matrase Reagent to release the cells from the extracellular matrix.

We observed a substantial, statistically significantly lower cell yield and cell viability in the final cell suspension isolated by just mechanical processing of human lipoaspirate without the use of Matrase Reagent, due to less efficient release of cells from the extracellular matrix when no enzyme was used. Specifically, the live cell yield of Transpose RT / Matrase isolated ADRCs was approximately twelve times higher in the final cell suspension than the live cell yield of Transpose RT / no Matrase isolated ADRCs. Of importance, the mean relative number of 86% viable Transpose RT / Matrase isolated ADRCs exceeded the proposed minimum threshold for viability of cells in the SVF of 70 % established by IFATS [71], whereas the mean relative number of 61% viable Transpose RT / no Matrase isolated ADRCs did not.

In Figure 16 cell yield and live cell yield data obtained in this study for Transpose RT / Matrase isolated ADRCs and Transpose RT / no Matrase isolated ADRCs are compared with corresponding data reported in the literature (c.f. Tables 1 and 2). For only six out of 39 enzymatic methods (15%) a higher cell yield was reported than for Transpose RT / Matrase isolated ADRCs found in this study (Fig 16A). However, the cell yield reported for one of these enzymatic methods (387×10^5^) [63] was approximately 80 times higher than the average cell yield reported for all other 38 methods listed in Table 1 (4.8×10^5^). Because the authors of [63] did not compare their extraordinarily high cell yield data with any of the data listed in Table 1, the cell yield reported in [63] should be treated with caution. Moreover, no cell viability data were provided in those six studies [57, 59, 60–63] that reported higher cell yield for enzymatic methods than found in this study for Transpose RT / Matrase isolated ADRCs. Accordingly, live cell yield could only be calculated for 25 out of the 39 enzymatic methods (64%) listed in Table 1, and live cell yield found in this study for Transpose RT / Matrase isolated ADRCs was higher than live cell yield calculated for any of these 25 enzymatic methods (Fig 16B). Likewise, live cell yield could only be calculated for one out of the ten enzymatic methods (10%) listed in Table 2, and live cell yield found in this study for Transpose RT / no Matrase isolated ADRCs was higher (Fig 16B).

**Figure 16.**
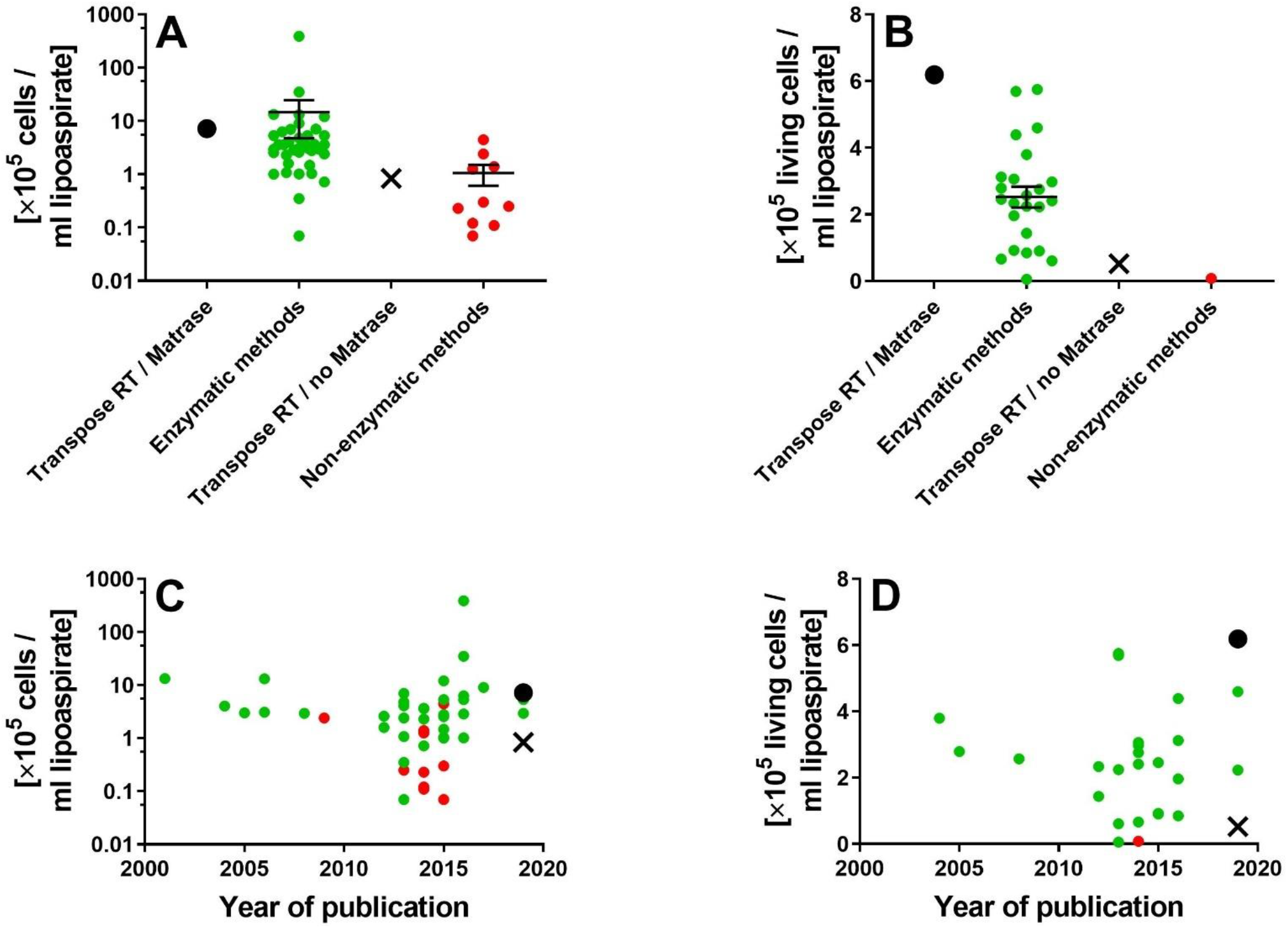
Cell yield and live cell yield obtained with Transpose RT / Matrase isolation of ADRCs and Transpose RT / no Matrase isolation of ADRCs compared with corresponding data reported in the literature. Panels A and B show individual data as well as mean ± standard error of the mean of cell yield (A) and live cell yield (B) obtained with Transpose RT / Matrase isolation of ADRCs (black dots), enzymatic methods for isolating ADRCs reported in the literature (listed in Table 1) (green dots), Transpose RT / no Matrase isolation of ADRCs (black crosses) and non-enzymatic methods for isolating ADRCs reported in the literature (listed in Table 2) (red dots). Panels C and D show the same data as a function of the year of publication.

For considerations of clinical usage, these findings have an important impact: the significantly lower viability of cells recovered just mechanically without Matrase Reagent means that nearly 40 % of the cells that would be transferred to a patient are not viable. In this regard is of note that Aronowitz and colleagues [43] already pointed out that the viability of the cells in the final cell suspension is a very important clinical factor because nonviable cells may provide no therapeutic value but could potentially lead to excess localized inflammation at the treatment site because of excessive cellular debris [43]. This should also be taken into account when considering the use of a non-enzymatic method for isolating ADRCs from human adipose tissue. As pointed out above, for nine out of the ten non-enzymatic methods for isolating ADRCs listed in Table 2 no cell viability data were provided, and both the non-enzymatic method described in [53] and Transpose RT / no Matrase isolation of ADRCs described in this study resulted in cell viability data that were below the threshold of 70% viable cells established by IFATS [71].

Cells isolated in this study from lipoaspirate were able to form CFUs independent of enzyme usage, indicating the presence of stem cells in both methods of isolating ADRCs from adipose tissue. However, Transpose RT / Matrase isolated ADRCs formed on average 16 times more CFUs per ml lipoaspirate than Transpose RT / no Matrase isolated ADRCs. These results are in line with previously reported studies comparing enzymatic to non-enzymatic extraction of ADRCs from lipoaspirate [56], as well as with results of cells recovered from adipose tissue or just from the tumescence fluid portion [60]. These data implicate that in order to even match the number of spheroids formed from Transpose RT / Matrase isolated ADRCs, 16 times more adipose tissue would be required as starting material for processing without enzyme. In other words, instead of 100 g patient derived adipose tissue for enzymatic processing, 1600 g for the just mechanical process would be required. This could mean a more complex recovery procedure and potentially higher morbidity, especially in patients at increased risk of bleeding. In this regard it was speculated in a recent study addressing treatment of osteoarthritis [74] that augmenting non-enzymatically isolated ADRCs with plateled rich plasma (PRP) could overcome the increased need for adipose tissue. However, no cell yield and no cell viability of the non-enzymatically isolated ADRCs were reported in [74]; no comparison of application of ADRCs with or without simultaneous application of PRP was performed; and clinical outcome was not validated with magentic resonance imaging, arthroscopy and microscopic evaluation of biopsies in [74].

Aside from substantial differences in cell yield, cell viability and live cell yield, Transpose RT / Matrase isolated ADRCs showed no statistically significant differences in the expression of regenerative cell-associated genes Oct4, Hes1 and Klf4 compared to Transpose RT / no Matrase isolated ADRCs. Besides this, both Transpose RT / Matrase isolated ADRCs and Transpose / no Matrase isolated ADRCs were able to differentiate into all three germ layers (i.e., into the adipogenic, osteogenic, hepatogenic and neurogenic lineages). This was in line with all enzymatic and non-enzymatic methods for isolating ADRCs listed in Tables 1 and 2 for which differentiation into all three germ layers was investigated.

In a joined position statement published by IFATS and the International Society for Cellular Therapy (ISCT) in 2013 regarding SVF and culture-expanded ASCs it was stated that primary stable positive surface markers for stromal cells are CD13, CD29, CD34 (>20%), CD44, CD73 and CD90 (>40%), whereas primary negative surface markers for stromal cells are CD31 (<20%) and CD45 (<50%) [71]. Furthermore, at least 20% of the SVF would contain a stromal cell population that is immunopositive for the surface marker CD34 and immunonegative for the surface markers CD31, CD45 and CD235a (i.e., CD31−/CD34+/CD45−/CD235a− cells) [71]. This statement was based on an earlier position statement published by ISCT in 2006 that described the following minimal criteria for defining multipotent mesenchymal stromal cells (MSCs): being adherent to plastic, expressing the surface markers CD73, CD90 and CD105, and having the ability to differentiate into osteoblasts, adipocytes and chondrocytes [75]. It should be pointed out that a major shortcoming of this definition of multipotent MSCs is the fact that, for example, fibroblasts are also adherent to plastic and express the surface markers CD73, CD90 and CD105, without having the ability to transdifferentiate into other lineages or being MSCs [72]. Furthermore, the true pluripotent stem cells do not yet express CD73, CD90 and CD105 [76]. Rather, expression of cell surface markers is a dynamic process. For example, when cultured in fetal bovine serum or platelet lysate culture media, MSCs can turn on new surface markers [76]. Alternatively, MSCs in culture can lose their surface marker expression, such as for example the loss of the previously expressed progenitor marker CD34 or the endothelial progenitor marker CD31 [76].

Nevertheless, Tables 6 and 7 summarize the relative amount of ADRCs expressing the surface markers CD13, CD29, CD34, CD44, CD73, CD90, CD31 and CD45 as reported in all studies describing enzymatic and non-enzymatic methods for isolating ADRCs listed in Tables 1 and 2 (note that in some studies surface markers were investigated but relative amounts of ADRCs expressing a certain surface marker or a combination of surface markers were not provided). The data summarized in Tables 6 and 7 demonstrate that (i) for only very few methods [46, 50, 51, 56] the relative amount of CD34^+^ ADRCs was determined, with substantial variation among methods (range, 35% - 81%); (ii) for most methods CD34 was determined together with at least one other surface marker, resulting in a range of published data between 0.8% (CD34^+^/CD90^−^/CD31^−^/CD45^−^/CD105^−^/CD146^+^ cells; [48]) and 44% (CD34^+^/CD31^−^/CD45^−^ cells; [40]); (iii) the relative amount of CD45^+^ ADRCs varied between 6% [57] and 50% [51] for enzymatic methods, and between 8% [57] and 82% [56] for non-enzymatic methods; (iv) for only few methods the relative amounts of CD13^+^ cells, CD29^+^ cells, CD44^+^ cells, CD73^+^ cells, CD90^+^ cells and CD31^+^ cells were determined; and (v) for no any method the relative amount of CD31^−^/CD34^+^/CD45^−^/CD235a^−^ cells (as proposed in [71]) was determined.

**Table 6.** Relative amount of ADRCs expressing certain surface markers as reported in studies describing non-enzymatic methods for isolating ADRCs.

| R | Y | CY | CD13 | CD29 | CD34 | CD44 | CD73 | CD90 | CD31 | CD45 |
| --- | --- | --- | --- | --- | --- | --- | --- | --- | --- | --- |
| [39] | 2015 | **0.0** | -- | -- | -- | -- | -- | -- | -- | -- |
| [40] | 2013 | **0.07** | -- | -- | 28<sup>(a)</sup> | -- | -- | -- | (a) | (a) |
| [40] | 2013 | **0.35** | -- | -- | 26<sup>(a)</sup> | -- | -- | -- | (a) | (a) |
| [41] | 2014 | **0.72** | -- | -- | -- | -- | -- | -- | -- | -- |
| [42] | 2015 | **1.00** | -- | -- | 20<sup>(b)</sup> | -- | -- | -- | 4<sup>(c)</sup> | -- |
| [43] | 2016 | **1.01** | -- | -- | 11<sup>(a)</sup> | -- | -- | -- | (a) | (a) |
| [42] | 2015 | **1.03** | -- | -- | 25<sup>(b)</sup> | -- | -- | -- | 5<sup>(c)</sup> | -- |
| [40] | 2013 | **1.07** | -- | -- | 22<sup>(a)</sup> | -- | -- | -- | (a) | (a) |
| [42] | 2015 | **1.47** | -- | -- | -- | -- | -- | -- | -- | -- |
| [44] | 2012 | **1.60** | -- | -- | 42<sup>(b)</sup> | -- | -- | -- | -- | 36 |
| [45] | 2014 | **2.30** | -- | -- | -- | -- | 60<sup>(d)</sup> | (d) | -- | (d) |
| [40] | 2013 | **2.41** | -- | -- | 44<sup>(a)</sup> | -- | -- | -- | (a) | (a) |
| [39] | 2015 | **2.55** | -- | -- | -- | -- | -- | -- | -- | -- |
| [44] | 2012 | **2.60** | -- | -- | 41<sup>(b)</sup> | -- | -- | -- | -- | 34 |
| [43] | 2016 | **2.85** | -- | -- | 9<sup>(a)</sup> | -- | -- | -- | (a) | (a) |
| [46] | 2019 | **2.93** | 33 | 76 | 60 | 60 | 48 | 68 | 45 | 23 |
| [47] | 2015 | **2.79** | -- | -- | 36<sup>(e)</sup> | -- | -- | (e) | -- | 5<sup>(f)</sup> |
| [48] | 2008 | **2.95** | -- | -- | 0.8<sup>(g)</sup> | -- | -- | -- | 8<sup>(h)</sup> | -- |
| [49] | 2005 | **3.00** | -- | -- | -- | -- | -- | -- | -- | -- |
| [50] | 2006 | **3.08** | 37 | 47 | 60 | 64 | 25 | 55 | 22 | -- |
| [51] | 2014 | **3.60** | -- | -- | 35 | -- | -- | -- | 8<sup>(i)</sup> | 50 |
| [52] | 2014 | **3.60** | -- | -- | -- | -- | -- | -- | -- | -- |
| [53] | 2014 | **3.60** | -- | -- | -- | -- | -- | -- | -- | -- |
| [54] | 2014 | **3.68** | -- | -- | -- | -- | -- | -- | -- | -- |
| [23] | 2004 | **4.04** | -- | -- | -- | -- | -- | -- | -- | -- |
| [55] | 2013 | **4.07** | -- | -- | -- | -- | -- | -- | -- | -- |
| [56] | 2013 | **4.80** | -- | 90 | 81 | 6 | 37 | 81 | -- | 28 |
| [57] | 2015 | **5.34** | -- | -- | 20<sup>(a)</sup> | -- | -- | -- | -- | 6 |
| [46] | 2019 | **5.35** | 48 | 89 | 67 | 68 | 59 | 76 | 55 | 31 |
| [43] | 2016 | **5.36** | -- | -- | 9<sup>(a)</sup> | -- | -- | -- | (a) | (a) |
| [43] | 2016 | **6.25** | -- | -- | 7<sup>(a)</sup> | -- | -- | -- | (a) | (a) |
| [58] | 2013 | **7.01** | -- | -- | 18<sup>(b)</sup> | -- | -- | -- | 27<sup>(i)</sup> | -- |
| [58] | 2013 | **7.02** | -- | -- | 43<sup>(b)</sup> | -- | -- | -- | 25<sup>(i)</sup> | -- |
| [59] | 2017 | **9.06** | -- | -- | -- | -- | -- | -- | -- | -- |
| [57] | 2015 | **12.1** | -- | -- | 19<sup>(a)</sup> | -- | -- | -- | -- | 13 |
| [60] | 2006 | **13.1** | -- | -- | 28<sup>(j)</sup> | -- | -- | -- | 4<sup>(k)</sup> | 30<sup>(l)</sup> |
| [61] | 2001 | **13.3** | -- | -- | -- | -- | -- | -- | -- | -- |
| [62] | 2016 | **35.0** | -- | -- | -- | -- | -- | -- | -- | -- |
| [63] | 2016 | **387** | -- | -- | -- | -- | -- | 22<sup>(m)</sup> | (m) | -- |

Surface marker data are given in percent. Abbreviations: R, reference; Y, year of publication; CY, cell yield [ $\times 10^5$ /ml lipoaspirate]; --, data not provided. (a), CD34<sup>+</sup>/CD31<sup>-</sup>/CD45<sup>-</sup> cells; (b), CD34<sup>+</sup>/CD31<sup>-</sup> cells; (c), CD34<sup>+</sup>/CD31<sup>+</sup> cells; (d), CD45<sup>-</sup>/CD73<sup>+</sup>/CD90<sup>+</sup> cells; (e), CD34<sup>+</sup>/CD90<sup>+</sup>/CD45<sup>-</sup>/CD146<sup>-</sup> cells; (f), CD34<sup>+</sup>/CD45<sup>+</sup>/CD14<sup>+</sup> cells; (g), CD34<sup>+</sup>/CD90<sup>+</sup>/CD31<sup>-</sup>/CD45<sup>-</sup>/CD105<sup>-</sup>/CD146<sup>+</sup> cells; (h), CD34<sup>+</sup>/CD90<sup>+</sup>/CD31<sup>+</sup>/CD45<sup>-</sup>/CD105<sup>-</sup>/CD146<sup>+</sup> cells; (i), CD34<sup>+</sup>/CD31<sup>+</sup> cells; (j) CD34<sup>+</sup>/CD90<sup>+</sup>/CD31<sup>-</sup>/CD45<sup>-</sup>/CD105<sup>-</sup>/CD146<sup>-</sup> cells; (k), CD34<sup>+</sup>/CD90<sup>+</sup>/CD31<sup>+</sup>/CD45<sup>-</sup>/CD105<sup>-</sup>/CD146<sup>+</sup> cells; (l), CD34<sup>-</sup>/CD31<sup>-</sup>/CD45<sup>-</sup>/CD105<sup>-</sup>/CD14<sup>-</sup>/CD15<sup>+</sup> cells; (m), CD90<sup>+</sup>/CD105<sup>+</sup>/CD31<sup>-</sup> cells.

**Table 7.** Relative amount of ADRCs expressing certain surface markers as reported in studies describing non-enzymatic methods for isolating ADRCs.

| R | Y | CY | CD13 | CD29 | CD34 | CD44 | CD73 | CD90 | CD31 | CD45 |
| --- | --- | --- | --- | --- | --- | --- | --- | --- | --- | --- |
| [39] | 2015 | **0.07** | -- | -- | -- | -- | -- | -- | -- | -- |
| [53] | 2014 | **0.11** | -- | -- | -- | -- | -- | -- | -- | -- |
| [45] | 2014 | **0.12** | -- | -- | -- | -- | 60<sup>(a)</sup> | (a) | -- | (a) |
| [45] | 2014 | **0.23** | -- | -- | -- | -- | 60<sup>(a)</sup> | (a) | -- | (a) |
| [56] | 2013 | **0.25** | -- | 48 | 24 | 5 | 9 | 23 | -- | 82 |
| [39] | 2015 | **0.30** | -- | -- | -- | -- | -- | -- | -- | -- |
| [64] | 2014 | **1.25** | -- | -- | -- | -- | -- | -- | -- | -- |
| [54] | 2014 | **1.39** | -- | -- | -- | -- | -- | -- | -- | -- |
| [65] | 2009 | **2.40** | -- | -- | -- | -- | -- | -- | -- | -- |
| [57] | 2015 | **4.44** | -- | -- | 25<sup>(a)</sup> | -- | -- | -- | -- | 8 |

Surface marker data are given in percent. Abbreviations: R, reference; Y, year of publication; CY, cell yield [ $\times 10^5$ /ml lipoaspirate]; --, data not provided. (a) CD45<sup>+</sup>/CD73<sup>+</sup>/CD90<sup>+</sup> cells; (b), CD34<sup>+</sup>/CD31<sup>+</sup>/CD45<sup>+</sup> cells.

Most importantly, the data summarized in Tables 6 and 7 do not show any correlation between the cell yield and any single surface marker or any combination of surface markers, respectively. Neither do these data allow any differentiation between reported enzymatic and non-enzymatic methods for isolating ADRCs from human adipose tissue. Rather, the vast majority of reported enzymatic and non-enzymatic methods were not characterized according to the position statements published by IFATS and ISCT [71, 75].

Considering the available data summarized in Tables 6 and 7 and the general concerns about characterizing ADRCs and MSCs by surface markers outlined above it appears reasonable to hypothesize that determining surface markers of ADRCs is in principle not suitable for characterizing a method for isolating ADRCs from adipose tissue. This was the reason why no such characterization was performed in this study.

In any case, the available surface marker data do not allow to conclude that using enzymes in the process of isolating ADRCs from human adipose tissue subjects the cells to substantial manipulation. Rather, the available surface marker data suggest that using enzymes in the process of isolating ADRCs from human adipose tissue does not subject the cells to substantial manipulation.

Of importance, after isolating cells with Matrase Reagent following the manufacturer’s instructions for use, the collagenase activity in the final cell suspension was not present (Fig 13). This suggests that the enzyme has only a supportive function in releasing the cells but has no presence or effect in the final cell suspension.

In summary, this study demonstrates that isolating ADRCs with the use of Matrase Reagent did not alter or induce biological characteristics, physiological functions or structural properties of the cells relevant for the intended use. Due to the high yield of viable, pluripotent cells in the final cell suspension isolated from lipoaspirate (or adipose tissue in general) with the Transpose RT system and the use of Matrase Reagent, cells are neither required to be cultured nor expanded, nor is any genetic manipulation necessary such as overexpression of embryonic genes as in the case of iPS cells.

Regarding the definition of minimal manipulation of cells, the U.S. Food and Drug Administration (FDA) has stated the following: “*Processing that does not alter the relevant biological characteristics of cells*” and “*Examples of relevant biological characteristics of cells or nonstructural tissues include differentiation and activation state, proliferation potential, and metabolic activity*” [77]. Furthermore, Transpose RT / Matrase isolation of ADRCs is performed directly at point of care, i.e., not *outside of a direct patient care setting* [78]. Recently, the FDA has approved clinical studies using Transpose RT / Matrase isolated ARDCs under Investigational Device Exemption (IDE) and not under Investigational New Drug Application (IND) [79–83].

The European Medicine Agency has defined advanced therapy medicinal products (ATMPs) as follows: “*Cells or tissues shall be considered ‘engineered’ if they fulfil at least one of the following conditions: 1) the cells or tissues have been subject to substantial manipulation, so that biological characteristics, physiological functions or properties relevant for the intended regeneration, repair or replacement are achieved. The manipulation listed in Annex I, in particular, shall not be considered as substantial manipulations; and 2) the cells or tissues are not intended to be used for the same essential function or functions in the recipient as in the donor*” [28]. Accordingly, in report EMA/CAT/228/2013 the European Medicine Agency has classified Transpose RT / Matrase isolated ARDCs as not substantially manipulated and classified as non-ATMP [84].

## CONCLUSIONS

For the practice of medicine our findings do not support the hypothesis that ADRCs should preferentially be isolated from adipose tissue without enzyme, as proposed in some studies in the literature [37, 74]. Rather, the results of this study as well as a wealth of data from the literature (Tables 1 and 2, Figs 1 and 16) clearly show that non-enzymatic (i.e., just mechanical) isolation of ADRCs from adipose tissue may result in significantly lower cell yield (and most probably also in lower cell viability) than enzymatic isolation of ADRCs from adipose tissue. In order to obtain a comparable cell yield just by mechanical processing without enzyme, larger initial amounts of adipose tissue must be recovered, let alone the potentially higher amount of non-living cells in the final cell suspension [43]. Transpose RT / Matrase isolated ADRCs are - upon the respective induction - capable to differentiate into all three germ layers without prior genetic manipulation. In line with earlier recommendations in the literature [43] characterization of ADRCs for clinical use should include analysis of nucleated cell count, nucleated cells per milliliter of tissue processed, cellular viability, level of residual enzymatic activity and data from CFU-F assay. On the other hand, analysis of surface markers of ADRCs should no longer be considered suitable for characterizing a method for isolating ADRCs from adipose tissue.

## REFERENCES

1. Forbes SJ, Rosenthal N. Preparing the ground for tissue regeneration: from mechanism to therapy. Nat Med. 2014; 20(8):857–69. doi: 10.1038/nm.3653 PMID: 25100531

2. Bianco P. “Mesenchymal” stem cells. Annu Rev Cell Dev Biol. 2014;30:677–704. doi: 10.1146/annurev-cellbio-100913-013132 PMID: 25150008

3. Goodell MA, Rando TA. Stem cells and healthy aging. Science 2015;350(6265):1199–204. doi: 10.1126/science.aab3388 PMID: 26785478

4. Nguyen PK, Rhee JW, Wu JC. Adult stem cell therapy and heart failure, 2000 to 2016: A systematic review. JAMA Cardiol. 2016;1(7):831–41. doi: 10.1001/jamacardio.2016.2225 PMID: 27557438

5. Cossu G, Birchall M, Brown T, De Coppi P, Culme-Seymour E, Gibbon S, et al. Lancet Commission: Stem cells and regenerative medicine. Lancet. 2018;391(10123):883–910. doi: 10.1016/S0140-6736(17)31366-1 PMID: 28987452

6. Schäffler A, Büchler C. Concise review: adipose tissue-derived stromal cells--basic and clinical implications for novel cell-based therapies. Stem Cells. 2007;25(4):818–27. doi: 10.1634/stemcells.2006-0589 PMID: 17420225

7. Alt E, Pinkernell K, Scharlau M, Coleman M, Fotuhi P, Nabzdyk C, et al. Effect of freshly isolated autologous tissue resident stromal cells on cardiac function and perfusion following acute myocardial infarction. Int J Cardiol. 2010; 144: 26–35. doi: 10.1016/j.ijcard.2009.03.124 PMID: 19443059

8. Klein SM, Vykoukal J, Li DP, Pan HL, Zeitler K, et al. Peripheral motor and sensory nerve conduction following transplantation of undifferentiated autologous adipose tissue-derived stem cells in a biodegradable U.S. Food and Drug administration-approved nerve conduit. Plast Reconstr Surg. 2016;138(1):132–9. doi: 10.1097/PRS.0000000000002291 PMID: 27348645

9. Konstantinow A, Arnold A, Djabali K, Kempf W, Gutermuth J, Fischer T, et al. Therapy of ulcus cruris of venous and mixed venous arterial origin with autologous, adult, native progenitor cells from subcutaneous adipose tissue: a prospective clinical pilot study. J Eur Acad Dermatol Venereol. 2017;31(12):2104–18. doi: 10.1111/jdv.14489 PMID: 28750144

10. Solakoglu Ö, Götz W, Kiessling MC, Alt C, Schmitz C, Alt EU. Improved guided bone regeneration by combined application of unmodified, fresh autologous adipose derived regenerative cells and plasma rich in growth factors: A first-in-human case report and literature review. World J Stem Cells. 2019;11(2):124–46. doi: 10.4252/wjsc.v11.i2.124 PMID: 30842809

11. McIntosh K, Zvonic S, Garrett S, Mitchell JB, Floyd ZE, Hammill L, et al. The immunogenicity of human adipose-derived cells: temporal changes in vitro. Stem Cells. 2006;24(5):1246–53. doi: 10.1634/stemcells.2005-0235 PMID: 16410391

12. Badimon L, Oñate B, Vilahur G. Adipose-derived mesenchymal stem cells and their reparative potential in ischemic heart disease. Rev Esp Cardiol (Engl Ed). 2015;68(7):599–611. doi: 10.1016/j.rec.2015.02.025 PMID: 26028258

13. Bora P, Majumdar AS. Adipose tissue-derived stromal vascular fraction in regenerative medicine: a brief review on biology and translation. Stem Cell Res Ther. 2017;8(1):145. doi: 10.1186/s13287-017-0598-y PMID: 28619097

14. Fujikawa T, Oh SH, Pi L, Hatch HM, Shupe T, Petersen BE. Teratoma formation leads to failure of treatment for type I diabetes using embryonic stem cell-derived insulin-producing cells. Am J Pathol. 2005; 166(6):1781–91. doi: 10.1016/S0002-9440(10)62488-1 PMID: 15920163

15. Swijnenburg RJ, Tanaka M, Vogel H, Baker J, Kofidis T, Gunawan F, et al. Embryonic stem cell immunogenicity increases upon differentiation after transplantation into ischemic myocardium. Circulation. 2005; 112(9 Suppl):I166–72. doi: 10.1161/CIRCULATIONAHA.104.525824 PMID: 16159810

16. King NM, Perrin J. Ethical issues in stem cell research and therapy. Stem Cell Res Ther. 2014; 5(4):85. doi: 10.1186/scrt474 PMID: 25157428

17. Ahmed RP, Ashraf M, Buccini S, Shujia J, Haider HKh. Cardiac tumorigenic potential of induced pluripotent stem cells in an immunocompetent host with myocardial infarction. Regen Med. 2011; 6(2):171–8. doi: 10.2217/rme.10.103 PMID: 21391851

18. Zhang Y, Wang D, Chen M, Yang B, Zhang F, Cao K. Intramyocardial transplantation of undifferentiated rat induced pluripotent stem cells causes tumorigenesis in the heart. PLoS One. 2011; 6(4):e19012. doi.: 10.1371/journal.pone.0019012 PMID 21552563

19. Trounson A. Potential pitfall of pluripotent stem cells. N Engl J Med. 2017; 377(5):490–1. doi: 10.1056/NEJMcibr1706906 PMID 28767348

20. Panés J, García-Olmo D, Van Assche G, Colombel JF, Reinisch W, Baumgart DC, et al. Expanded allogeneic adipose-derived mesenchymal stem cells (Cx601) for complex perianal fistulas in Crohn’s disease: a phase 3 randomised, double-blind controlled trial. Lancet. 2016; 388(10051):1281–90. doi: 10.1016/S0140-6736(16)31203-X PMID: 27477896

21. Álvaro-Gracia JM, Jover JA, García-Vicuña R, Carreño L, Alonso A, Marsal S, et al. Intravenous administration of expanded allogeneic adipose-derived mesenchymal stem cells in refractory rheumatoid arthritis (Cx611): results of a multicentre, dose escalation, randomised, single-blind, placebo-controlled phase Ib/IIa clinical trial. Ann Rheum Dis. 2017; 76(1):196–202. doi: 10.1136/annrheumdis-2015-208918 PMID: 27269294

22. Kastrup J, Haack-Sørensen M, Juhl M, Harary Søndergaard R, Follin B, Drozd Lund L, et al. Cryopreserved off-the-shelf allogeneic adipose-derived stromal cells for therapy in patients with ischemic heart disease and heart failure-a safety study. Stem Cells Transl Med. 2017;6:1963–71. doi: 10.1002/sctm.17-0040 PMID: 28880460

23. Aust L, Devlin B, Foster SJ, Halvorsen YD, Hicok K, du Laney T, et al. Yield of human adipose-derived adult stem cells from liposuction aspirates. Cytotherapy. 2004; 6(1):7–14. doi: 10.1080/14653240310004539 PMID: 14985162

24. Yu H, Lu K, Zhu J, Wang J. Stem cell therapy for ischemic heart diseases. Br Med Bull. 2017; 121(1):135–54. doi: 10.1093/bmb/ldw059 PMID: 28164211

25. Safwani WK, Makpol S, Sathapan S, Chua KH. Alteration of gene expression levels during osteogenic induction of human adipose derived stem cells in long-term culture. Cell Tissue Bank. 2013;14(2):289–301. doi: 10.1007/s10561-012-9309-1 PMID: 22476937

26. Pan Q, Fouraschen SM, de Ruiter PE, Dinjens WN, Kwekkeboom J, Tilanus HW, et al. Detection of spontaneous tumorigenic transformation during culture expansion of human mesenchymal stromal cells. Exp Biol Med. 2014;239(1):105–15. doi: 10.1177/1535370213506802 PMID: 24227633

27. Jiang T, Xu G, Wang Q, Yang L, Zheng L, Zhao J, et al. In vitro expansion impaired the stemness of early passage mesenchymal stem cells for treatment of cartilage defects. Cell Death Dis. 2017;8(6):e2851. doi: 10.1038/cddis.2017.215. PMID: 28569773

28. European Medicines Agency. Scientific recommendation on classification of advanced therapy medicinal products. Article 17 – Regulation (EC) No 1394/2007, 2012.

29. Mitchell JB, McIntosh K, Zvonic S, Garrett S, Floyd ZE, Kloster A, et al. Immunophenotype of human adipose-derived cells: temporal changes in stromal-associated and stem cell-associated markers. Stem Cells. 2006;24(2):376–85. doi: 10.1634/stemcells.2005-0234. PMID: 16322640

30. El-Badawy A, Amer M, Abdelbaset R, Sherif SN, Abo-Elela M, Ghallab YH, et al. Adipose stem cells display higher regenerative capacities and more adaptable electro-kinetic properties compared to bone marrow-derived mesenchymal stromal cells. Sci Rep. 2016;6:37801. doi: 10.1038/srep37801. DOI: 27883074

31. Toyserkani NM, Jørgensen MG, Tabatabaeifar S, Jensen CH, Sheikh SP, Sørensen JA. Concise review: a safety assessment of adipose-derived cell therapy in clinical trials: a systematic review of reported adverse events. Stem Cells Transl Med. 2017;6(9):1786–94. doi: 10.1002/sctm.17-0031. PMID: 28722289

32. Sheu JJ, Lee MS, Wallace CG, Chen KH, Sung PH, Chua S, et al. Therapeutic effects of adipose derived fresh stromal vascular fraction-containing stem cells versus cultured adipose derived mesenchymal stem cells on rescuing heart function in rat after acute myocardial infarction. Am J Transl Res. 2019;11(1):67–86. PMID: 30787970

33. Nyberg E, Farris A, O’Sullivan A, Rodriguez R, Grayson WL. Comparison of SVF and passaged ASCs as point-of-care agents for bone regeneration. Tissue Eng Part A. 2019 Feb 8: Epub ahead of print. doi: 10.1089/ten.TEA.2018.0341. PMID: 30734661

34. You D, Jang MJ, Kim BH, Song G, Lee C, Suh N, et al. Comparative study of autologous stromal vascular fraction and adipose-derived stem cells for erectile function recovery in a rat model of cavernous nerve injury. Stem Cells Transl Med. 2015;4(4):351–8. doi: 10.5966/sctm.2014-0161. PMID: 25792486

35. Aronowitz JA, Lockhart RA, Hakakian CS. Mechanical versus enzymatic isolation of stromal vascular fraction cells from adipose tissue. Springerplus. 2015; 4:713. doi: 10.1186/s40064-015-1509-2 PMID: 26636001

36. Oberbauer E, Steffenhagen C, Wurzer C, Gabriel C, Redl H, Wolbank S. Enzymatic and non-enzymatic isolation systems for adipose tissue-derived cells: current state of the art. Cell Regen. 2015; 4:7. doi: 10.1186/s13619-015-0020-0 PMID: 26435835

37. Condé-Green A, Kotamarti VS, Sherman LS, Keith JD, Lee ES, Granick MS, et al. Shift toward mechanical isolation of adipose-derived stromal vascular fraction: review of upcoming techniques. Plast Reconstr Surg Glob Open. 2016; 4(9):e1017. doi: 10.1097/GOX.0000000000001017 PMID: 27757339

38. van Dongen JA, Tuin AJ, Spiekman M, Jansma J, van der Lei B, Harmsen MC. Comparison of intraoperative procedures for isolation of clinical grade stromal vascular fraction for regenerative purposes: a systematic review. J Tissue Eng Regen Med. 2018; 12(1):e261–74. doi: 10.1002/term.2407 PMID: 28084666

39. Gentile P, Scioli MG, Orlandi A, Cervelli V. Breast reconstruction with enhanced stromal vascular fraction fat grafting: What is the best method? Plast Reconstr Surg Glob Open. 2015; 3(6):e406. doi: 10.1097/GOX.0000000000000285 PMID: 26180707

40. Aronowitz JA, Ellenhorn JD. Adipose stromal vascular fraction isolation: a head-to-head comparison of four commercial cell separation systems. Plast Reconstr Surg. 2013; 132(6):932e–39e. doi: 10.1097/PRS.0b013e3182a80652 PMID: 24281640

41. Gotoh M, Yamamoto T, Kato M, Majima T, Toriyama K, Kamei Y, et al. Regenerative treatment of male stress urinary incontinence by periurethral injection of autologous adipose-derived regenerative cells: 1-year outcomes in 11 patients. Int J Urol. 2014; 21(3):294–300. doi: 10.1111/iju.12266 PMID: 24033774

42. SundarRaj S, Deshmukh A, Priya N, Krishnan VS, Cherat M, Majumdar AS. Development of a system and method for automated isolation of stromal vascular fraction from adipose tissue lipoaspirate. Stem Cells Int. 2015; 2015:109353. doi: 10.1155/2015/109353 PMID: 26167182

43. Aronowitz JA, Lockhart RA, Hakakian CS, Birnbaum ZE. Adipose stromal vascular fraction isolation: A head-to-head comparison of 4 cell separation systems #2. Ann Plast Surg. 2016; 77(3):354–62. doi: 10.1097/SAP.0000000000000831 PMID: 27220016

44. Güven S, Karagianni M, Schwalbe M, Schreiner S, Farhadi J, Bula S, et al. Validation of an automated procedure to isolate human adipose tissue-derived cells by using the Sepax® technology. Tissue Eng Part C Methods. 2012; 18(8):575–82. doi: 10.1089/ten.TEC.2011.0617 PMID: 22372873

45. Condé-Green A, Rodriguez RL, Slezak S, Singh DP, Goldberg NH, McLenithan J. Comparison between stromal vascular cells’ isolation with enzymatic digestion and mechanical processing of aspirated adipose tissue. Plast Reconstr Surg. 2014; 134(4S-1):54 doi: 10.1097/01.prs.0000455394.06800.62

46. Nürnberger S, Lindner C, Maier J, Strohmeier K, Wurzer C, Slezak P, et al. Adipose-tissue-derived therapeutic cells in their natural environment as an autologous cell therapy strategy: the microtissue-stromal vascular fraction. Eur Cell Mater. 2019; 37:113–33. doi: 10.22203/eCM.v037a08 PMID: 30793275

47. Granel B, Daumas A, Jouve E, Harlé JR, Nguyen PS, Chabannon C, et al. Safety, tolerability and potential efficacy of injection of autologous adipose-derived stromal vascular fraction in the fingers of patients with systemic sclerosis: an open-label phase I trial. Ann Rheum Dis. 2015; 74(12):2175–82. doi: 10.1136/annrheumdis-2014-205681 PMID: 25114060

48. Lin K, Matsubara Y, Masuda Y, Togashi K, Ohno T, Tamura T, et al. Characterization of adipose tissue-derived cells isolated with the Celution system. Cytotherapy. 2008; 10(4):417–26. doi: 10.1080/14653240801982979 PMID: 18574774

49. Kakudo N, Tanaka Y, Morimoto N, Ogawa T, Kushida S, Hara T, et al. Adipose-derived regenerative cell (ADRC)-enriched fat grafting: optimal cell concentration and effects on grafted fat characteristics. J Transl Med. 2013; 11:254. doi: 10.1186/1479-5876-11-254 PMID: 24107489

50. Mitchell JB, McIntosh K, Zvonic S, Garrett S, Floyd ZE, Kloster A, et al. Immunophenotype of human adipose-derived cells: temporal changes in stromal-associated and stem cell-associated markers. Stem Cells. 2006; 24(2):376–85. doi: 10.1634/stemcells.2005-0234 PMID: 16322640

51. Fraser JK, Hicok KC, Shanahan R, Zhu M, Miller S, Arm DM. The Celution® system: Automated processing of adipose-derived regenerative cells in a functionally closed system. Adv Wound Care. 2014; 3(1):38–45. doi: 10.1089/wound.2012.0408 PMID: 24761343

52. Dos-Anjos Vilaboa S, Navarro-Palou M, Llull R. Age influence on stromal vascular fraction cell yield obtained from human lipoaspirates. Cytotherapy. 2014; 16(8):1092–7. doi: 10.1016/j.jcyt.2014.02.007 PMID: 24726656

53. Markarian CF, Frey GZ, Silveira MD, Chem EM, Milani AR, Ely PB, et al. Isolation of adipose-derived stem cells: a comparison among different methods. Biotechnol Lett. 2014; 36(4):693–702. doi: 10.1007/s10529-013-1425-x PMID: 24322777

54. Millan A, Landerholm T, Chapman JR. Comparison between collagenase adipose digestion and Stromacell mechanical dissociation for mesenchymal stem cell separation. McNair Scholars J CSUS 2014; 15:86–101

55. Muscari C, Bonafè F, Fiumana E, Oranges CM, Pinto V, Caldarera CM, et al. Comparison between stem cells harvested from wet and dry lipoaspirates. Connect Tissue Res. 2013; 54(1):34–40. doi: 10.3109/03008207.2012.717130 PMID: 22853627

56. Shah FS, Wu X, Dietrich M, Rood J, Gimble JM. A non-enzymatic method for isolating human adipose tissue-derived stromal stem cells. Cytotherapy. 2013; 15(8):979–85. doi: 10.1016/j.jcyt.2013.04.001 PMID: 23725689

57. Domenis R, Lazzaro L, Calabrese S, Mangoni D, Gallelli A, Bourkoula E, et al. Adipose tissue derived stem cells: in vitro and in vivo analysis of a standard and three commercially available cell-assisted lipotransfer techniques. Stem Cell Res Ther. 2015; 6:2. doi: 10.1186/scrt536 PMID: 25559708

58. Doi K, Tanaka S, Iida H, Eto H, Kato H, Aoi N, et al. Stromal vascular fraction isolated from lipo-aspirates using an automated processing system: bench and bed analysis. J Tissue Eng Regen Med. 2013; 7(11):864–70. doi: 10.1002/term.1478 PMID: 22438241

59. Raposio E, Bertozzi N. How to isolate a ready-to-use adipose-derived stem cells pellet for clinical application. Eur Rev Med Pharmacol Sci. 2017; 21(18):4252–60 PMID: 29028071

60. Yoshimura K, Shigeura T, Matsumoto D, Sato T, Takaki Y, Aiba-Kojima E, et al. Characterization of freshly isolated and cultured cells derived from the fatty and fluid portions of liposuction aspirates. J Cell Physiol. 2006; 208(1):64–76. doi: 10.1002/jcp.20636 PMID: 16557516

61. Zuk PA, Zhu M, Mizuno H, Huang J, Futrell JW, Katz AJ, et al. Multilineage cells from human adipose tissue: implications for cell-based therapies. Tissue Eng 2001; 7(2):211–28. doi: 10.1089/107632701300062859 PMID: 11304456

62. Raposio E, Caruana G, Petrella M, Bonomini S, Grieco MP. A standardized method of isolating adipose-derived stem cells for clinical applications. Ann Plast Surg. 2016; 76(1):124–6. doi: 10.1097/SAP.0000000000000609 PMID: 26418805

63. Grasys J, Kim BS, Pallua N. Content of soluble factors and characteristics of stromal vascular fraction cells in lipoaspirates from different subcutaneous adipose tissue depots. Aesthet Surg J. 2016; 36(7):831–41. doi: 10.1093/asj/sjw022 PMID: 26906346

64. Raposio E, Caruana G, Bonomini S, Libondi G. A novel and effective strategy for the isolation of adipose-derived stem cells: minimally manipulated adipose-derived stem cells for more rapid and safe stem cell therapy. Plast Reconstr Surg. 2014; 133(6):1406–9. doi: 10.1097/PRS.0000000000000170 PMID: 24867723

65. Baptista LS, do Amaral RJ, Carias RB, Aniceto M, Claudio-da-Silva C, Borojevic R. An alternative method for the isolation of mesenchymal stromal cells derived from lipoaspirate samples. Cytotherapy. 2009; 11(6):706–15. doi: 10.3109/14653240902981144 PMID: 19878057

66. Astori G, Vignati F, Bardelli S, Tubio M, Gola M, Albertini V, et al. “In vitro” and multicolor phenotypic characterization of cell subpopulations identified in fresh human adipose tissue stromal vascular fraction and in the derived mesenchymal stem cells. J Transl Med. 2007; 5:55. doi: 10.1186/1479-5876-5-55 PMID: 17974012

67. Brickman JM, Serup P. Properties of embryoid bodies. Wiley Interdiscip Rev Dev Biol. 2017; 6(2). doi: 10.1002/wdev.259. doi: 10.1002/wdev.259 PMID: 27911036

68. Izant JG, McIntosh JR. Microtubule-associated proteins: a monoclonal antibody to MAP2 binds to differentiated neurons. Proc Natl Acad Sci USA. 1980; 77:741–5. doi: 10.1073/pnas.77.8.4741 PMID: 7001466

69. Lee MK, Tuttle JB, Rebhun LI, Cleveland DW, Frankfurter A. The expression and posttranslational modification of a neuron-specific beta-tubulin isotype during chick embryogenesis. Cell Motil Cytoskeleton 1990; 17: 118–32. doi: 10.1002/cm.970170207 PMID: 2257630

70. Bhang SH, Kwon SH, Lee S, Kim GC, Han AM, Kwon YH, Kim BS. Enhanced neuronal differentiation of pheochromocytoma 12 cells on polydopamine-modified surface. Biochem Biophys Res Commun. 2013; 430: 1294–300. doi: 10.1016/j.bbrc.2012.11.123 PMID: 23261471

71. Bourin P, Bunnell BA, Casteilla L, Dominici M, Katz AJ, March KL, et al. Stromal cells from the adipose tissue-derived stromal vascular fraction and culture expanded adipose tissue-derived stromal/stem cells: a joint statement of the International Federation for Adipose Therapeutics and Science (IFATS) and the International Society for Cellular Therapy (ISCT). Cytotherapy. 2013; 15(6):641–648. doi: 10.1016/j.jcyt.2013.02.006 PMID: 23570660

72. Alt E, Yan Y, Gehmert S, Song YH, Altman A, Gehmert S, et al. Fibroblasts share mesenchymal phenotypes with stem cells, but lack their differentiation and colony-forming potential. Biol Cell. 2011; 103(4):197–208. doi: 10.1042/BC20100117 PMID: 21332447

73. Kageyama R, Ohtsuka T, Kobayashi T. The Hes gene family: repressors and oscillators that orchestrate embryogenesis. Development. 2007; 134(7):1243–51. doi: 10.1242/dev.000786 PMID: 17329370

74. Słynarski K, Baszczeski F, Lipinski L. Treatment of osteoarthritis – adipose derived stem cell & PRP therapy. Sportärztezeitung. 2017; 3: 14–19

75. Dominici M, Le Blanc K, Mueller I, Slaper-Cortenbach I, Marini F, Krause D, et al. Minimal criteria for defining multipotent mesenchymal stromal cells. The International Society for Cellular Therapy position statement. Cytotherapy. 2006; 8(4):315e7. doi: 10.1080/14653240600855905 PMID: 16923606

76. Alt E, Schmitz C, Bai X. Why and how patients’ own adult stem cells are the next generation of medicine. Preprints 2019, 2019040200. doi: 10.20944/preprints201904.0200.v1

77. U.S. Department of Health and Human Services. Regulatory considerations for human cells, tissues, and cellular and tissue-based products: minimal manipulation and homologous use. Guidance for industry and Food and Drug Administration staff. Published December 2017. Available at https://www.fda.gov/ucm/groups/fdagov-public/@fdagov-bio-gen/documents/document/ucm585403.pdf (accessed on June 26th, 2019)

78. U.S. Department of Health and Human Services. Evaluation of Devices Used with Regenerative Medicine Advanced Therapies. Guidance for Industry. Published February 2019. Available at https://www.fda.gov/media/120266/download (accessed on June 26th, 2019)

79. Hurd J. Safety and efficacy adipose-derived stem cell injection partial thickness rotator cuff tears. IDE 16956. ClinicalTrials.gov Identifier: NCT02918136. Available at https://www.clinicaltrials.gov/ct2/show/NCT02918136 (accessed on June 26th, 2019)

80. Coots BK. healing chronic venous stasis wounds with autologous cell therapy. IDE 17214. ClinicalTrials.gov Identifier: NCT02961699. Available at https://www.clinicaltrials.gov/ct2/show/NCT02961699 (accessed on June 26th, 2019)

81. Boetel T. Safety of adipose-derived regenerative cells injection for treatment of osteoarthritis of the facet joint. IDE 17991. ClinicalTrials.gov Identifier: NCT03513731. Available at https://www.clinicaltrials.gov/ct2/show/NCT03513731 (accessed on June 26th, 2019)

82. Hurd J. Autologous adult adipose-derived regenerative cell injection into chronic partial-thickness rotator cuff tears. ClinicalTrials.gov Identifier: NCT03752827. Available at https://www.clinicaltrials.gov/ct2/show/NCT03752827 (accessed on June 26th, 2019)

83. Vandermark R. Healing osteoarthritic joints in the wrist with adult ADRCs. IDE 17984. ClinicalTrials.gov Identifier: NCT03503305. Available at https://www.clinicaltrials.gov/ct2/show/NCT03503305 (accessed on June 26th, 2019)

84. European Medicines Agency. Scientific recommendation on classification of advanced therapy medicinal products. Article 17 – Regulation (EC) No 1394/2007. EMA/CAT/228/2013, 2013

